# Mutations in legume genes that influence symbiosis create a complex selective landscape for rhizobial symbionts

**DOI:** 10.1101/2024.09.11.612462

**Authors:** Sohini Guha, Regina B. Bledsoe, Jeremy Sutherland, Brendan Epstein, Gwendolyn M. Fry, Vikram Venugopal, Siva Sankari, Alejandra Gil Polo, Garrett Levin, Barney Geddes, Nevin D. Young, Peter Tiffin, Liana T. Burghardt

## Abstract

In the mutualism between leguminous plants and rhizobia bacteria, rhizobia live inside root nodules, creating potential for host genes to shape the rhizobial selective environment. Many host genes that affect symbiosis have been identified; however, the extent to which these genes affect selection acting on rhizobia is unknown. In this study, we inoculated 18 *Medicago truncatula* symbiotic mutants (including mutants that alter Nodule Cysteine-Rich (NCR) peptide production, plant defense, and nodule number regulation) with a mixture of 86 *Sinorhizobium meliloti* strains. Most mutations resulted in reduced host benefits, but the effects on rhizobial benefit (i.e., relative strain fitness) varied widely, revealing widespread host-by-strain fitness interactions. Genome-wide association analyses identified variants on rhizobial replicons pSymA and pSymB as important in mediating strain fitness responses to host mutations. Whereas most top variants affected rhizobial fitness with one host mutant (limited effect variants), nine affected fitness across six or more host mutations. These pervasive variants occurred primarily on pSymA, the symbiotic replicon, and include *fixL* and some metabolic genes. In contrast to the limited effect variants, variants with pervasive positive effects on strain fitness when host genes were mutated tended to adversely affect fitness in wild-type hosts. Competition assays across *Medicago* genotypes confirmed a pervasive role for one candidate (malonyl-CoA synthase), and AlphaFold multimer modelling suggests that many rhizobial top candidates could interact with host NCR peptides. Our results reveal how host genetic variation affects rhizobial fitness, setting the stage for improving rhizobial inoculants and breeding legume hosts better adapted to multi-strain environments.

## Introduction

Many bacteria live in close association with eukaryotic hosts, and a subset of these bacterial taxa significantly influence host fitness^1^. If host genetic variation alters selective pressures on bacteria, that variation can shape bacterial evolution. These genome x genome (GxG) interactions are an essential prerequisite for co-evolution, and a potential mechanism for maintaining genetic variation in bacterial and host populations^2^. Most examples of (GxG) interaction involve pathogens, including a host immune receptor and microbial targets (e.g., human HLA and HIV epitopes^3^) or plant R genes^4^. However, examinations of quantitative pathogen resistance leveraging techniques such as co-GWAS (Genome Wide Association Studies)^5,6^ suggest complex genetic architectures underlie these host-pathogen interactions^7,8^. More recently, studies of (GxG) interactions have expanded to mutualisms^9–12^. These studies suggest that (GxG) interactions in mutualisms may be equally, if not more, genetically complex than in parasitism^13–18^. To what extent and in what ways host genes alter bacterial selection has important implications for bacterial evolution and deploying beneficial microbes in translational contexts such as agriculture^19,20^. Here, we take a novel approach to measuring (GxG) interactions by measuring how over a dozen host mutations alter the selective landscape experienced by scores of strains of mutualistic bacteria.

Legume hosts and rhizobium bacteria are a tractable system for studying how individual host genes alter the selection of their microbial mutualists. Rhizobia inhabit diverse environments, including soil, the rhizosphere, and root nodules, where they experience selection pressures that shape their fitness^21^. In nodules, rhizobia convert atmospheric nitrogen into plant assimilable forms and receive photosynthates for growth^22,23^. Host genetic variation can influence rhizobial fitness within root nodules^16,24–26^ in ways likely shaped by nodule development and subsequent nitrogen fixation^27^. Decades of work have resulted in extensive collections of mutant lines in model legumes, including *Lotus* and *Medicago*, which have been exhaustively used to dissect the functional genetics of nodule formation and function. Most research on these mutants has focused on their effect on host benefits (e.g., shoot biomass) and investment (e.g., nodule number/size). Far less attention has been paid to the effect of host mutations on rhizobia^28^. However, doing so is necessary to clarify the potential for host genes to shape the rhizobial fitness.

Measuring the effect of host genes on rhizobia requires considering three aspects of rhizobial fitness—absolute fitness, relative fitness, and nodule strain diversity. Because most nodules are initiated by a single rhizobium, rhizobial population sizes in nodules indicate the rewards an individual bacterium receives from forming a nodule^28,29^. However, measuring population size alone does not enable an estimate of fitness difference among strains. Assessing strain-level relative fitness requires comparing strain frequencies before and after exposure to a selective environment, such as host nodules. A Select and Resequence (S&R) approach leverages whole-genome shotgun (WGS) sequencing to enable high-throughput tracking of strain frequencies^25,30^. This method allows for the direct quantification of selection pressures on rhizobia, revealing how selective an environment is (nodule strain diversity) and which strains are enriched or depleted compared to the initial community (relative strain fitness)^25,30^. A key advantage of the S&R method is that, because all strains are already sequenced, GWAS can be used to identify genetic variants associated with rhizobial relative fitness^16,25,30,31^.

Work in the legume rhizobium mutualism suggests that host genetic variation can alter these metrics of rhizobial fitness^16,24,25,30^. For instance, host mutations that increase plant defence responses (*nad1*) reduce the number of rhizobia released from each nodule, whereas disrupting genes that cause hosts to form more nodules (*sunn1, sunn4, rdn1*) can increase rhizobia released at the plant level^32^. Other host genes like *rj2* and *rj4* in soybean restrict the diversity of strains that form nodules^33,34^. Disruption of the *Sinorhizobium* gene encoding HRRP peptidase, potentially involved in processing host NCR peptides, can restrict the rhizobial strains in nodules by modulating host discrimination or defence thresholds^35^. Lastly, disruptions within a host gene can alter *relative strain fitness.* For instance, members of the NPD gene family in *Medicago truncatula* differentially influenced which strain frequencies increased in nodules relative to the inoculated strain community^30^. Taken together, these studies highlight the potential for single host genes to reshape the selective environment and alter the evolution of rhizobia.

Here, we characterize how genetically disrupting host genes alters selection on their nitrogen-fixing rhizobium symbionts *in planta*. To accomplish this goal, we chose eighteen existing *Medicago truncatula* mutant lines with disruptions in the known symbiotic genes. These genes affect a wide range of host phenotypes and encompass the developmental trajectory of nodulation. We inoculated these mutants with a mixture of 86 *Sinorhizobium meliloti* strains (Fig. 1). We studied standard host investment and benefit traits, as well as multiple aspects of rhizobial fitness (Table 1). We then used a S&R approach to measure the frequencies of rhizobial strains (relative strain fitness) in nodules^25^ and followed up with GWAS, to identify variants in rhizobium genes that altered strain fitness when host genes were perturbed. We examined the following questions: 1) Which host symbiosis genes alter rhizobial fitness? 2) Do altered host genotypes have strain-specific effects on rhizobial fitness (i.e., is there evidence of genome x genome interactions)? 3) If yes, which rhizobium genes harbour genetic variation underlying the observed shifts in strain fitness, and are any of those variants selected in multiple host mutations? And 4) Are any host investment or benefit traits correlated with rhizobial fitness traits? Whereas few host genotypes altered rhizobial population sizes or nodule strain diversity, many hosts did shift relative strain fitness. Most rhizobial genes associated with strain fitness shifts were found to be important within specific hosts, although eighteen variants, (primarily on pSymA) were important with six or more hosts, suggesting candidate genes that may play fundamental roles in in-plant rhizobial fitness.

**Fig. 1:**
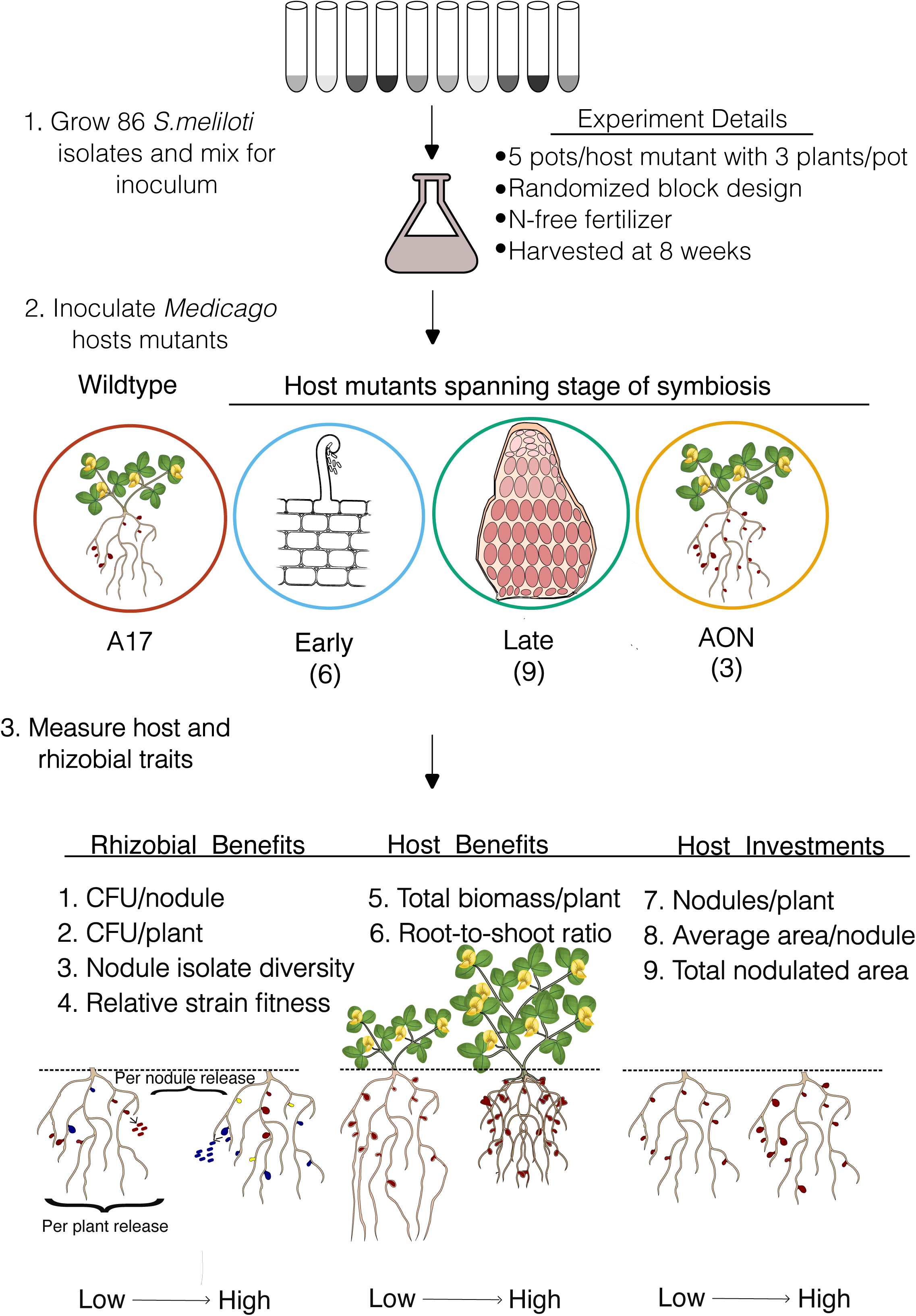
Schematic overview of the experiment used to evaluate rhizobial fitness and host traits across different host mutant backgrounds. A set of 86 *Sinorhizobium* strains was cultured and combined into a mixed inoculum. This inoculum was applied to *Medicago truncatula* wild-type (A17) and host mutants with gene disruptions that act at different stages of symbiosis: Early, Late, and Autoregulation of Nodulation (AON). The number of host mutants per stage is indicated in parentheses. Host and rhizobial traits were measured at the nodule and whole-plant scales. The experimental conditions are clarified under ‘Experimental Details’. Each treatment included five pots per host mutant with three plants per pot, grown in randomized blocks under nitrogen (N)-free conditions for 8 weeks. Traits measured include indicators of rhizobial benefit, host benefit, and host investment (Refer to Table 1 for more details). Different coloured nodules indicate nodules formed by different rhizobial strains.

**Table 1:**
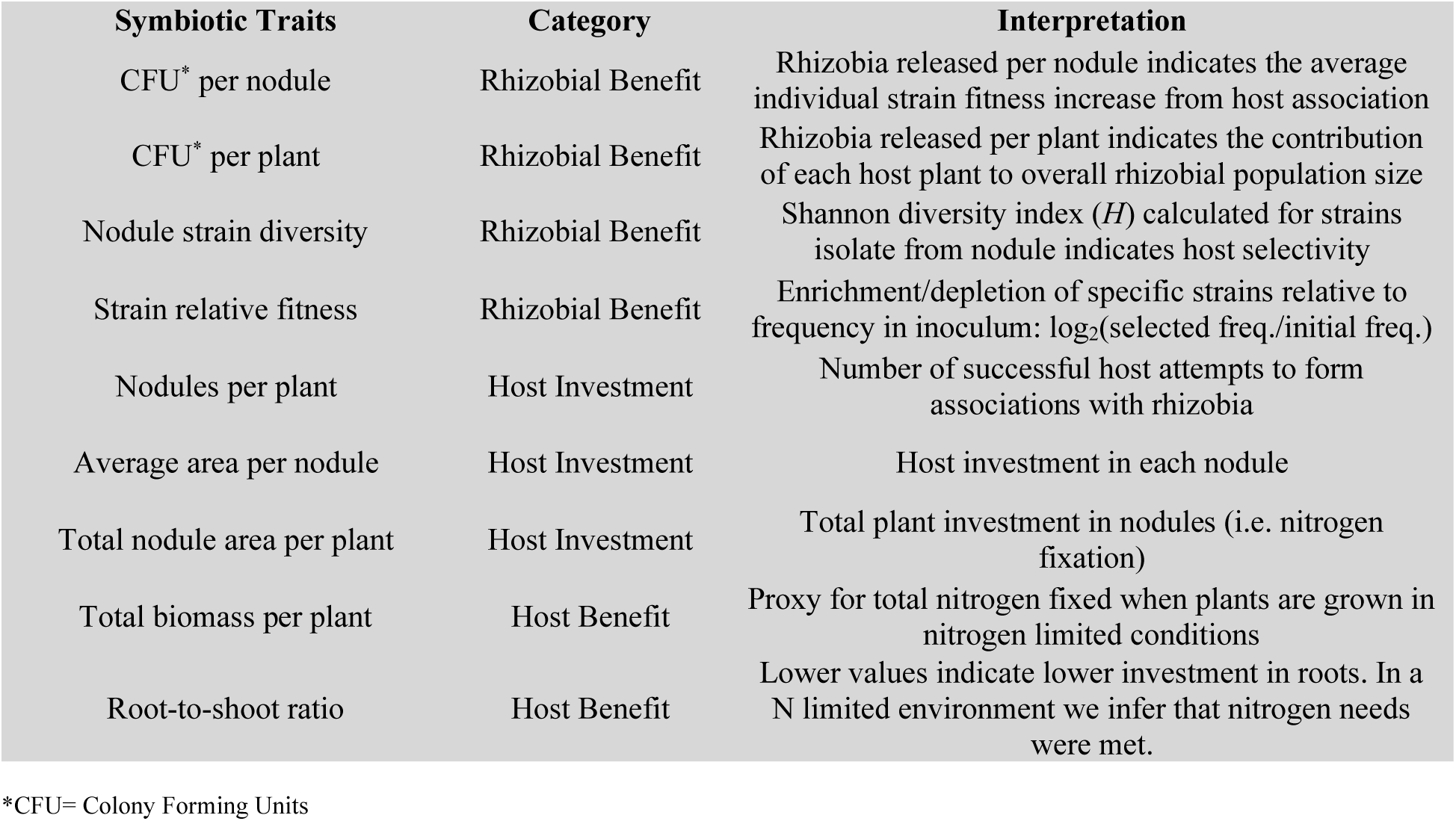
Definitions of the symbiotic traits examined.

## Materials and methods

### Host symbiotic mutants

The development of symbiosis in nodules is complex, with each step regulated by multiple hosts and rhizobium genes^27,36,37^. To encompass these steps, we chose 18 *M. truncatula* symbiosis mutants (Table S1), which we grouped into three categories: early-acting, late-acting, and regulation of nodule number (Autoregulation of nodule number; AON)^38^. The early-acting genes (*hcl, ipd3*, *latd, dmi1-2*, *dmi2-3,* and *dmi3-1*) affect the initial processes, such as Nod Factor recognition and nodule organogenesis^39–43^. The late-acting genes (*nad1, dnf1, dnf2, dnf4, dnf2/dnf1, dnf5/dnf2, dnf6, dnf7*) influence nitrogen fixation^32,44^, whereas the regulation of nodule number genes are in the AON (Autoregulation Of Nodulation) pathway (*sunn1, sunn4, rdn1*)^45,46^. Whereas these categorizations help set expectations for the impact of the host mutations, each gene may have additional pleiotropic effects. Together, these mutants provide a broad view of the effects of host symbiosis genes on their rhizobial partners in multi-strain inoculations. Seeds for all host mutant lines were obtained from Dr. Dong Wang, Department of Biochemistry and Molecular Biology, University of Massachusetts, Amherst. The host mutant lines used here were generated through forward genetic screens and thus may carry additional background mutations. Our aim is not to assign strict causality to individual host mutations but rather to examine how alterations in the host genetic landscape broadly influence rhizobial fitness.

### Initial inoculum preparation

We created a mixed inoculum using 86 sequenced *Sinorhizobium meliloti* (henceforth *S. meliloti*) strains, which were part of a collection isolated from soils from 21 locations in Spain and France, using the nodules of *Medicago* plants as a “trap” for capturing *S. meliloti* isolates^47^. We grew each strain separately for 72 hours in Tryptone-Yeast extract (TY) media at 28°C, shaking at 120 rpm, before mixing equal volumes of each culture to create an initial inoculum with roughly equal proportions of each strain. To inoculate plants, we added 60 ml of the multi-strain mixture to 6 litres of sterile 0.85% NaCl solution and applied 10^7^ cells per pot. This density was chosen to reflect *S. meliloti* populations in field soil^48^ and to avoid triggering quorum-sensing mechanisms^49^, which typically occur at higher cell densities. We measured rhizobial density in the inoculum by plating a dilution series and counting the number of colonies. We measured the strain frequencies in the inoculum by centrifuging eight 1 ml aliquots at 18,000 x g for 2 minutes, removing the supernatant, and then freezing the pellet at – 20°C before extracting DNA. Initial communities were sequenced, and frequencies inferred (Fig. S1) as outlined in the Rhizobial Fitness section below.

### Greenhouse experiment to measure multi-strain symbiotic phenotypes

We grew five replicate pots of each of the 18 *Medicago* mutants plus wildtype A17 (Fig. 1). Before planting, seeds were scarified by gently abrading with sandpaper, surface sterilized by soaking in a 10% bleach solution for 90 seconds and rinsed in de-ionized water. Seeds were stratified on wet filter paper at 4°C in the dark for 2 days, then planted in pots containing steam-sterilized Sunshine Mix #5. Germinated seedlings were thinned to three plants per pot. All pots were inoculated 48 hours after seed planting with the initial 86-strain inoculum. Following inoculation, pots were randomized into five spatial blocks with one representative of each genotype per block. Plants were watered as needed and fertilized weekly using increasing concentrations of N-free Fårhaeus medium^50^. Uninoculated plants were included to monitor cross-contamination. Pots were harvested and traits measured 8 weeks post-inoculation. We measured nodule number per plant, average area per nodule and total nodulated area per plant as measures of host investment, total biomass per plant and root-to-shoot ratio as measures of host benefit, and population size of rhizobia released per plant and per nodule, nodule strain diversity and relative strain fitness in nodule pools as measures of rhizobial fitness. After 8 weeks, we harvested inoculated plants and pooled nodules from ∼ 3 plants per pot to ensure enough nodules (>100 /pot) to limit stochastic variation and saturate diversity. None of the uninoculated plants formed nodules. We photographed nodule pools per replicate to assess nodule number and size using ImageJ. Plant biomass was measured by weighing the plant roots and shoots after oven-drying at 75 °C for 3 days. Because not all plants survived in every pot, the number of plants occasionally varied. To control for this variation, all host traits were normalized per plant for analysis.

### Rhizobial fitness

We surface-sterilized the nodules from each pot in 10% bleach for 1 minute, rinsed the nodules in sterile water, and homogenized them for 90 seconds in 4 ml sterile 0.85% sodium chloride solution using an Omni TH tissue homogenizer. We estimated the number of rhizobia released from the nodule pools by dilution plating a 500-1000 ul aliquot of the homogenate and counting the number of colonies formed. We then estimated the ***absolute fitness*** of the rhizobia on both a per-plant and per-nodule level by dividing the total number of colonies by the number of plants or nodules harvested from each pot and plotting the log_10_ transformation of this value. The remaining nodule homogenate was spun at 400 x g for 10 min at 10°C. to pellet the bacteroids and the plant debris. We collected the supernatant enriched for undifferentiated rhizobial cells, pelleted them at 18000 x g for 5 min, pipetted off the supernatant before freezing the pellets at –20 °C until DNA extraction. DNA was extracted using a Qiagen DNeasy® Plant extraction kit following the manufacturer’s protocol, except we used 3 μL of RNase in the first step and 50 μL of nuclease water for the final elution.

Samples were sequenced using Illumina short-read technology at the Huck Genomic Core Facility at The Pennsylvania State University. Indexed libraries were generated using the Illumina DNA Prep Kit. 12X libraries were pooled and sequenced on a 150 x 150 paired-end NextSeq High Output run, resulting in an average read depth of ∼75. We trimmed the reads using TrimGalore! (v0.4.1, www.bioinformatics.babraham.ac.uk/projects/trim_galore/) with default settings, except with minimum read length: 100 bp; quality threshold: 30 (−-quality); and minimum adapter match: 3 (−stringency), used bwa mem (v0.7.17) with default settings to align reads to the *S. meliloti* USDA1106 genome (NCBI BioProject: PRJNA388336) and estimated strain frequencies using HARP^51^. Strain frequencies were used to 1) estimate ***nodule strain diversity*** using Shannon’s H index (via the diversity function from the vegan package in R) and 2) calculate ***relative strain fitness*** of each strain in each replicate of a particular host mutant by taking the log_2_ of the ratio of the final strain frequency in the nodules to the initial frequency in the inoculum (median initial frequency = 0.0092, 5%-95% quantile: 0.006-0.018). The log transformation puts increases and decreases in proportional representation on the same scale (e.g., a fourfold decrease and increase will have a value of −2 and 2, respectively). Strains with frequency estimates of 0 in the selected communities were assigned a fitness value of −8, less than the lowest measurable reductions in other strains^25^.

### Statistical analyses

To assess the effects of the host mutants and the symbiotic stage on plant traits and rhizobial fitness, we used linear models and analysis of variance (ANOVA) for the univariate traits and redundancy analysis (RDA) for the multivariate trait of relative strain fitness. All analyses were conducted in the R statistical environment (version 4.3.2). For each univariate trait, we fit two linear models using the `lm` () function: one that includes host genotype and block, and a second that includes the symbiotic disruption stage and block as additive predictors. We compared variation explained by the two models using the adjusted *R*^2^ (`anova()`function) to infer whether models that included mutant identity explained more variance than the symbiotic stage alone. Because the genotypes are nested within the stage of symbiosis, we ran two separate linear models to avoid introducing collinearity. We then compared the variance explained by each model. To ensure this comparison was not biased by differences in the number of levels (degrees of freedom) in each predictor, we used the adjusted *R*², which corrects for model complexity and sample size. This approach allows us to assess the relative explanatory power of broad functional categories (stage) versus specific genetic backgrounds (genotypes) on rhizobial community structure. We log10-transformed average nodule area/plant and total nodule area traits to reduce heteroscedasticity of variance. To assess pairwise differences between A17 and each mutant for each trait, we used Tukey’s Honest Significant Difference (HSD) tests. Comparisons with an adjusted *P*-value ***<*** 0.05 were considered statistically significant. To investigate how host genotype or symbiotic stage altered the relative fitness of rhizobial strains (a multivariate trait), we performed an RDA (`rda()` function of the vegan package, v2.6-4) using relative strain fitness as the response matrix and host genotype or symbiotic stage as the explanatory variable. The significance of each model was assessed by extracting the adjusted *R*^2^ and via permutation-based ANOVA (1,000 permutations) using the `anova()` function with `by = “terms”`. To test whether the relative strain fitness within each mutant differed from A17, we ran the model for each mutant-A17 pairwise contrast. Predictors with *P* < 0.05 indicated significant changes in strain composition.

### Genome-wide association study on relative strain fitness shifts in mutants vs. wildtype nodules

The relative strain fitness shift of each strain for each host mutant was calculated by subtracting the relative strain fitness in A17 from the relative strain fitness in the host mutants. We conducted 15 GWAS (three mutants formed no nodules) to identify rhizobial variants underlying host mutation-dependent ***strain fitness shifts*** for each nodulating mutant. To be thorough, we also conducted a GWAS on strain fitness in each host genotype individually, which highlights variants associated with strain success in host nodules generally, as well as the effect of the perturbations. For each of these GWAS, we ran a linear mixed model (LMM) using GEMMA (version 0.98.3). Analyses were run separately for the three replicons (chromosome, pSymA, pSymB) to account for the different population structures. We used a published database of 30,076 SNPs representing previously defined linkage-disequilibrium (LD) groups within *S. meliloti*^16^. SNP filtering was performed with bcftools-1.18^52^ and PLINK 1.9^53^ based on minor allele frequency (0.05) and missingness (0.2). We selected the top five variant groups for each mutation-driven fitness shift tested based on their *P*-value ranks, cross-compared in the other 15 GWAS. To control for multiple tests while enabling comparisons across replicons that differed in variant group size from ∼ 3,000 to 12,000, we adopted a Bonferroni-corrected *P*-value cutoff of *P* < 10^-^⁵ to approximate a nominal *P* < 0.05 across replicons. This choice of a consistent intermediate threshold aligns with our goal of examining selection landscapes rather than identifying individual causal variants. This allowed us to categorise a rhizobial candidate as ‘pervasive’ if it ranked in the top ten within six or more host mutants or ‘limited effect’ if it appeared in five or fewer. We focused on ranks rather than *P*-values for this comparative analysis, prioritizing variants with the most evidence, regardless of differences in power or genetic architecture across host genetic perturbations.

### Functional validation of malonyl CoA synthase gene through competition assays

To validate the functional role of a candidate gene identified through GWAS, we mutated the malonyl-CoA synthase gene within the genetically amenable *S. meliloti* Rm1021 background (mutant strain named RmND1439). We competed the mutant against the corresponding wild type Rm1021, across four host genotypes (A17, *dnf3*, *dnf7*, and *sunn4*). The mutant and wild-type strains were cultured individually and mixed in equal proportions to create the initial inoculum. Strain representation in the inoculum and post-inoculation nodule pools was quantified by performing a dilution series and counting colony-forming units (CFU) on selective and non-selective plates.

Competitive fitness was expressed as the log₂ fold change in the proportion of RmND1439 recovered from the nodules relative to its proportion in the initial inoculum. Detailed methods for strain construction and competition assays are provided in the Supplementary Methods.

### In-silico prediction of the rhizobial candidate gene functions through protein-protein interaction using AlphaFold-Multimer and Foldseek

To obtain additional information about the potential for the protein products of the candidate rhizobial genes to interact with host proteins such as the NCR peptides directly, we performed in silico protein–protein interaction predictions using AlphaFold-Multimer^54,55^. The rhizobial candidate proteins were putatively annotated using Foldseek^56^. To infer candidate gene involvement during symbiosis, we also listed the nodule-enrichment scores of each rhizobial candidate from the Symbimics database (https://iant.toulouse.inra.fr/symbimics/) database^57^. These scores quantify the likelihood of the rhizobial candidate gene involvement during root nodule symbiosis, with higher values indicating stronger nodule-specific gene expression. Detailed procedures for structural modelling, interaction scoring, and data filtering are provided in the Supplementary Methods.

## Results

### Effect of symbiotic mutations on rhizobial population sizes

Community inoculation resulted in nodulation across all host mutants except for *dmi1-2*, *dmi2-3, and dmi3-1,* consistent with the expectation that these genes are required for nodule formation (Fig. 2A). For the 15 mutants that formed nodules, we measured the number of rhizobia released from the nodules and tested if each mutant differed significantly from A17 plants. For instance, *ipd3* releases ∼ 2.1×10^5^ rhizobial cells per nodule and ∼2.435×10^7^ per plant, a 10-fold decrease compared to A17 (Fig. 2B, C). For both, rhizobia released per nodule and plant, symbiosis stage did not explain nearly as much variation in rhizobia population sizes as individual host mutations (Table S2). Only two mutants significantly reduced the rhizobial population size at both per plant and nodule levels: *ipd3 (*per plant *P=*0.0001; per nodule *P*=0.04) and *nad1 (*per plant *P* = 0.02; per nodule *P*=0.03), and no mutations increased population size (Table S3). Because *ipd3* and *nad1* do not produce significantly different numbers of nodules compared to A17 (*P* > 0.05), the smaller per-plant rhizobial population size is primarily due to reduced reproduction of rhizobia within nodules. This makes sense given that *nad1* host mutant is known to evoke a defense-like response in nodules that kills reproductively active bacteria, and *ipd3* host mutant has aborted infection threads, the part of the nodule in which reproductively competent rhizobia are found^43,58^.

**Fig. 2:**
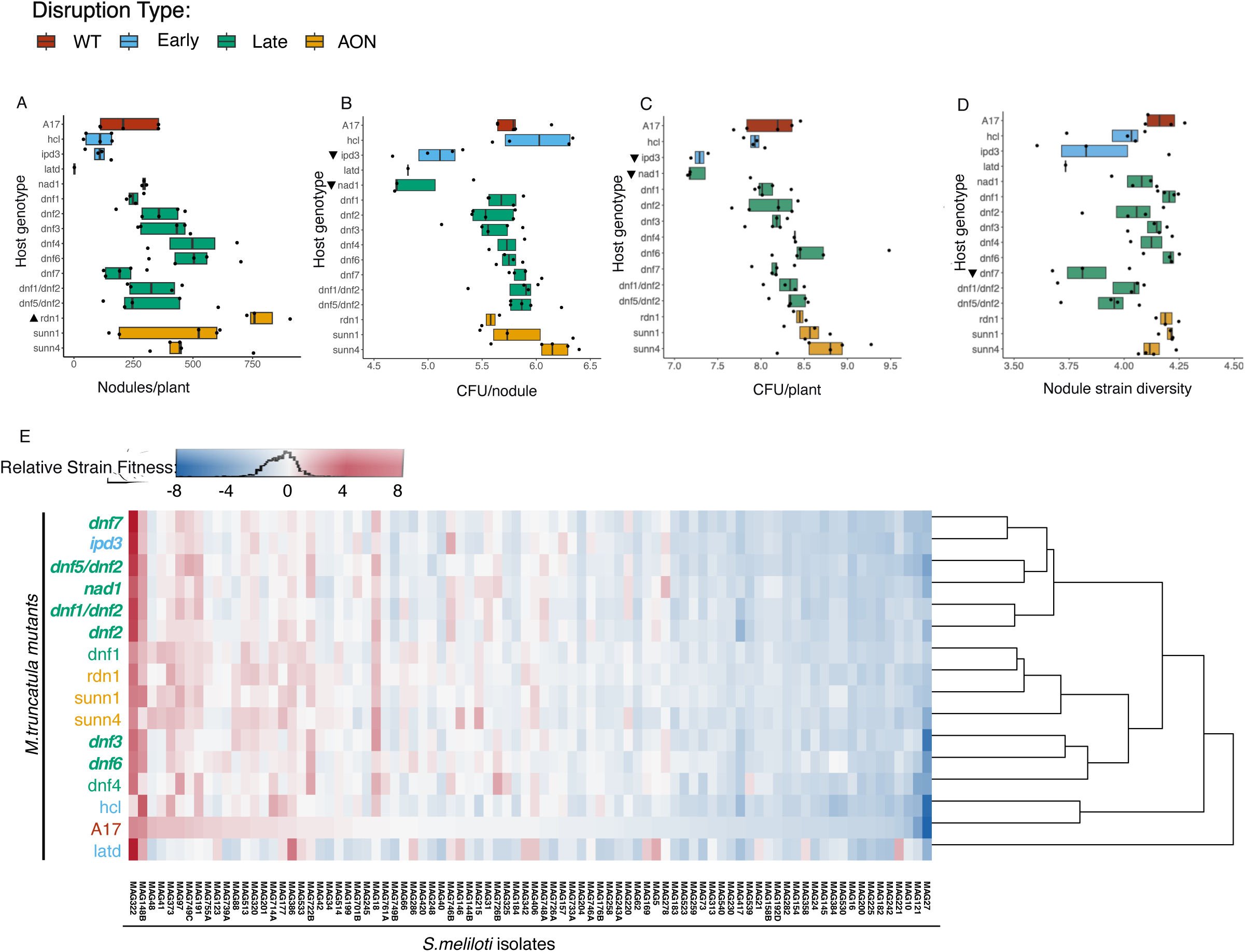
Host symbiotic mutations influence rhizobial fitness. (A) Host mutations altered nodule numbers in consistent and predictable ways. (B–C) Host genotype also influenced rhizobial fitness, as measured by log₁₀-transformed colony-forming units per nodule and per plant (CFU/nodule) (B) and CFU /plant (C). (D) Nodule strain diversity was largely unaffected by host mutations. Closed triangles indicate a significant increase or decrease relative to wild-type A17 (based on Tukey-adjusted *P* values). Boxplots are color-coded by the symbiotic stage disrupted by each host mutation (see Table S1). (E) Heatmap showing the median relative fitness of each of 86 *S. meliloti* strains across the different host genotypes. Strains enriched relative to their initial inoculum appear in shades of red; depleted strains appear in shades of blue. Host genotypes are hierarchically clustered based on similarity in strain fitness profiles. Strains are ranked by their relative fitness in A17. Mutants in bold differ significantly from A17 (see Table S5 for RDA paired with the PERMANOVA results).

### Host symbiotic genes shape relative strain success

The total rhizobial population released from individual nodules and plants does not tell us whether the effects of the mutations are strain specific. We examined strain-specific effects of the mutants in two ways. First, we evaluated the *nodule strain diversity,* as measured by Shannon’s H, in each of the 15 nodulating mutants compared to A17. This analysis revealed that *dnf7* was the only mutant with significantly reduced *nodule strain diversity* (*P* = 0.009, Fig. 2C, Table S3). Second, we compared the relative fitness of each strain in the nodules of each host mutant to A17 (Fig. 2E). Hierarchical clustering revealed that strain relative fitness did not cluster solely by symbiotic stage and that the *latd* nodule strain fitness was quite different from all other hosts, including wildtype. The *latd* host mutant results should be interpreted carefully— *latd* uniquely had a single estimate for rhizobial fitness traits, because plants formed so few nodules that we had to combine nodules across all pots to create a large enough sample for sequencing. A redundancy analysis with PERMANOVAs reinforced this finding. Whereas the symbiosis stage explained some variation in relative strain fitness, symbiotic genes had strong individual effects (Fig. S2, Table S4, S5). Both early-acting (*ipd3* and *nad1)* and late-acting (all *dnf* mutants except *dnf4,* which was marginal at *P* = 0.06, and *dnf1*) significantly altered rhizobial relative fitness in nodules (Fig. 2E). The *dnf* mutants did not cluster together in the RDA space, reinforcing that defects in nitrogen fixation did not produce uniform effects on rhizobial enrichment. Some strains consistently exhibited high or low fitness across all host genotypes (e.g., MAG322 vs. MAG27), but the fitness of most strains depended on host symbiotic mutations and often changed the rank order of relative strain fitness of the strains compared to wildtype (Fig. S3)., Strain shifts were often specific to individual host mutants; for example, MAG726B, MAG701B, and MAG201 had high fitness in *dnf4* but low fitness in *latd*. The host-dependent changes in strain fitness allow us to use GWAS to identify rhizobial allelic variants associated with the observed strain fitness shifts caused by each mutation relative to wildtype.

### Rhizobial genes underlying strain fitness shifts in response to host mutations were primarily located on extrachromosomal replicons

To determine genetic variants associated with shifts in strain fitness when the host genes were disrupted, we performed 16 GWAS, one for the shifts in strain fitness caused by each of the 15 mutants (relative fitness in mutant-relative fitness in A17) and for comparison, one for relative fitness in A17 (Fig.3, Fig. S4). We found 9 linkage groups on pSymA and 17 linkage groups on pSymB that were strongly associated with strain fitness shifts (*P <* 10^−5^) (Table 2, Table S6). In contrast, linkage groups on the chromosome had weaker effects (none with *P <* 10^−5^), except for LD group 838 (*P=* 6.38 x 10^−6^), which was associated with increased strain fitness with *dnf6* and *latd*. The *P* values of variants across different replicons may differ in part due to replicon-specific patterns of relatedness among strains. However, the impact of population structure on GWAS power is not always straightforward to predict. For instance, the chromosome may exhibit long-range linkage disequilibrium, resulting in large linkage blocks that tag multiple genes, which can both help or hinder association resolution. Additionally, we ran a second GWAS to identify the genes underlying strain fitness in each host (Fig. S5, Table S7). As expected, within the top five candidates, 11% were shared between A17 and the host mutants (Table S8), the rest were unique for the mutants, indicating host genetic make-up influences strain fitness (Table S9). Whereas there was some overlap in genetic variants across the two GWAS (Table S10), the candidates for shifts in strain fitness in response to host genetic perturbations were largely distinct (only 20 out of 113 variant groups of strain fitness shifts overlapped with strain fitness) (Table S11). Below, we focus on candidates from the fitness shift analyses because we are interested in rhizobial responses to changes in the host environment, rather than general fitness determinants.

**Fig. 3:**
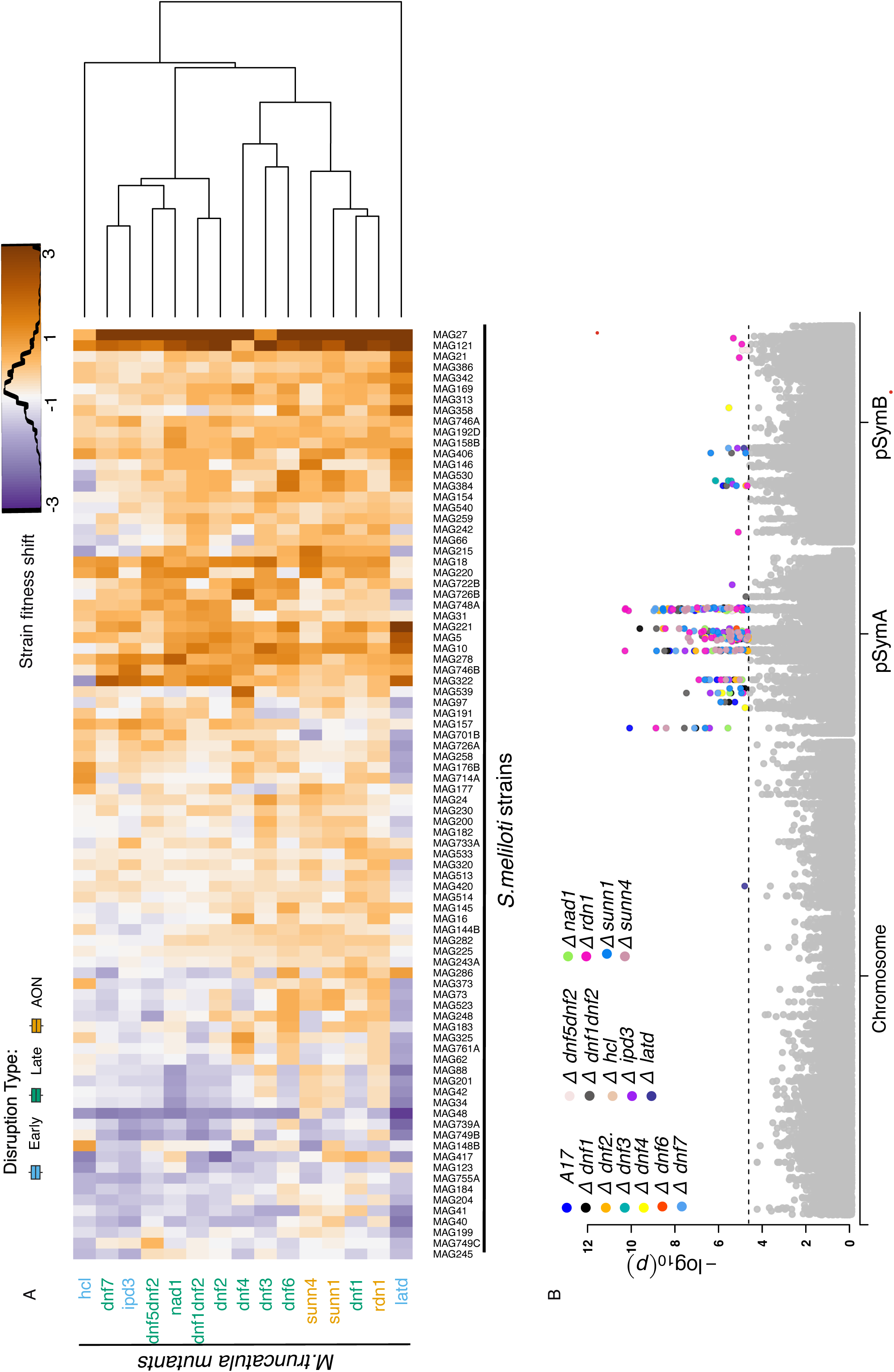
GWAS of strain fitness shifts in host mutants. (A) The rhizobial strain fitness shifts for each of the 86 strains, calculated by subtracting individual relative strain fitness measured in A17 from the relative strain fitness in the host mutants. The heatmap is hierarchically based on the similarity of median ‘shift’ values across hosts. Shades of orange denote positive fitness shift compared to A17, whereas shades of purple denote negative fitness shift compared to A17. (B) Manhattan plot indicating the SNP (single-nucleotide polymorphism) positions, based on S. *meliloti* USDA1106 association with the relative strain fitness shifts across the 15 host mutants and strain fitness in A17. The dotted line indicates the *P-*value threshold used for identifying candidate genes (*p* <10^^−5^), and significant associations are color-coded according to host mutation and A17. The GWAS was run separately on each replicon to account for different population structures. The ‘Δ’ indicates ‘strain fitness shift’ in that host mutant. For comparison, we also ran the GWAS on A17 relative strain fitness.

**Table 2:**
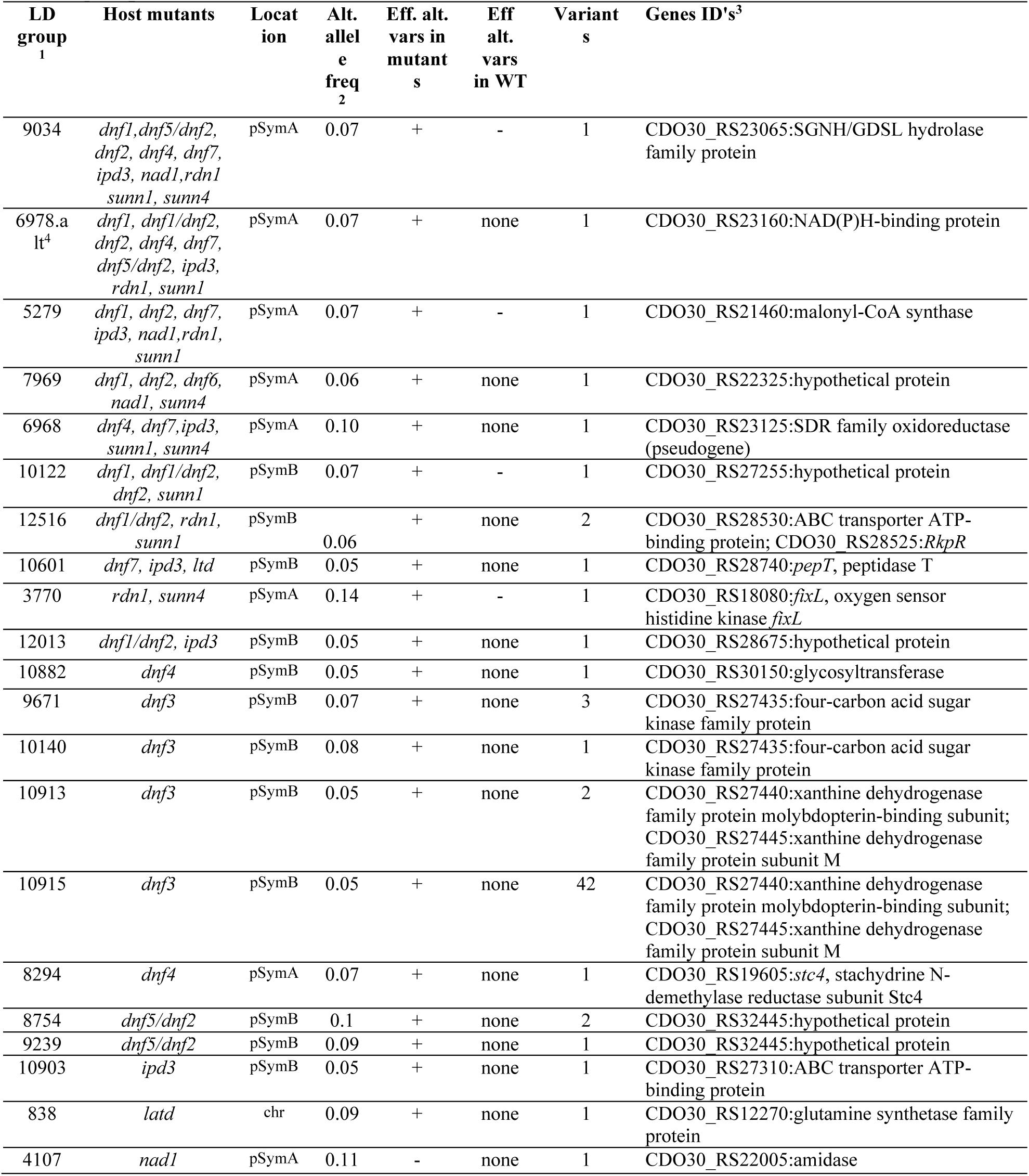
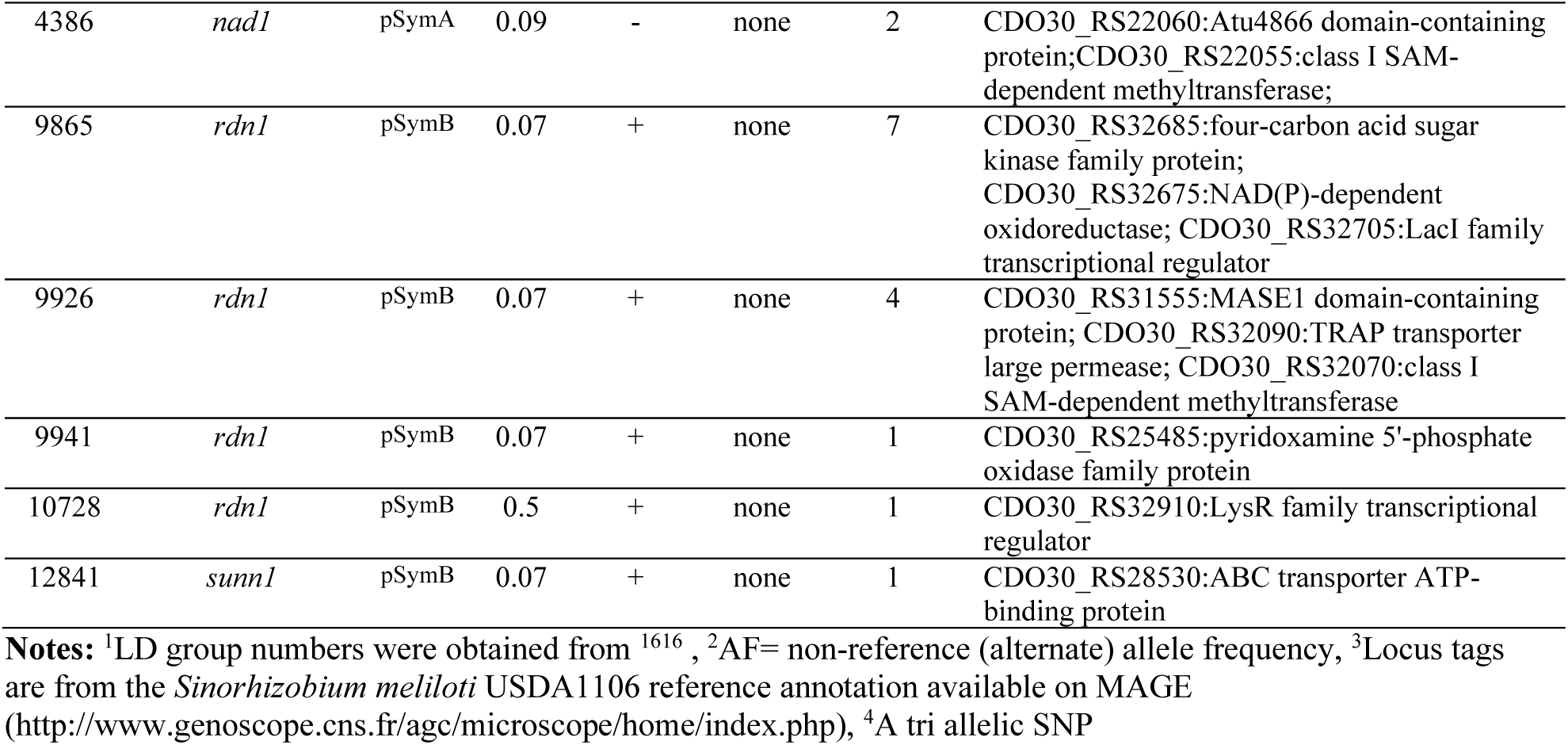
Candidate rhizobial genes in small variant groups (three genes or fewer) associated with (*P* < 10^-5^) variation in rhizobial relative strain fitness. See Table S6 for information on all variant groups.

### GWAS identifies rhizobial genes with pervasive and specific effects on strain fitness shifts

To study the genomic landscape underlying rhizobial fitness shifts in response to host mutations, we identified the top five variants for each host mutant (in total, 114 variant groups) based on *P*-ranks (Table S6). Then we compared the *P*-value ranks and effect sizes of these groups across all the host mutants (Fig. 4, Fig. S5, Table S12). Most of the candidates on the chromosome were classified as ‘limited effect’ variant groups because they ranked among the top ten candidates, in five or fewer host mutations (Table S12). Exceptions included a massive chromosomal variant group (901) with 144 variants in 65 genes, among the top ten candidates among nine host mutants. Six variants with pervasive effects were found on pSymB. On pSymB, four small variant groups were top-ranked in at least three host mutants (ABC transporter and RkpR, a dihydrodipicolinate synthase family protein, peptidase T, and a hypothetical protein (Fig. S6).

**Fig. 4:**
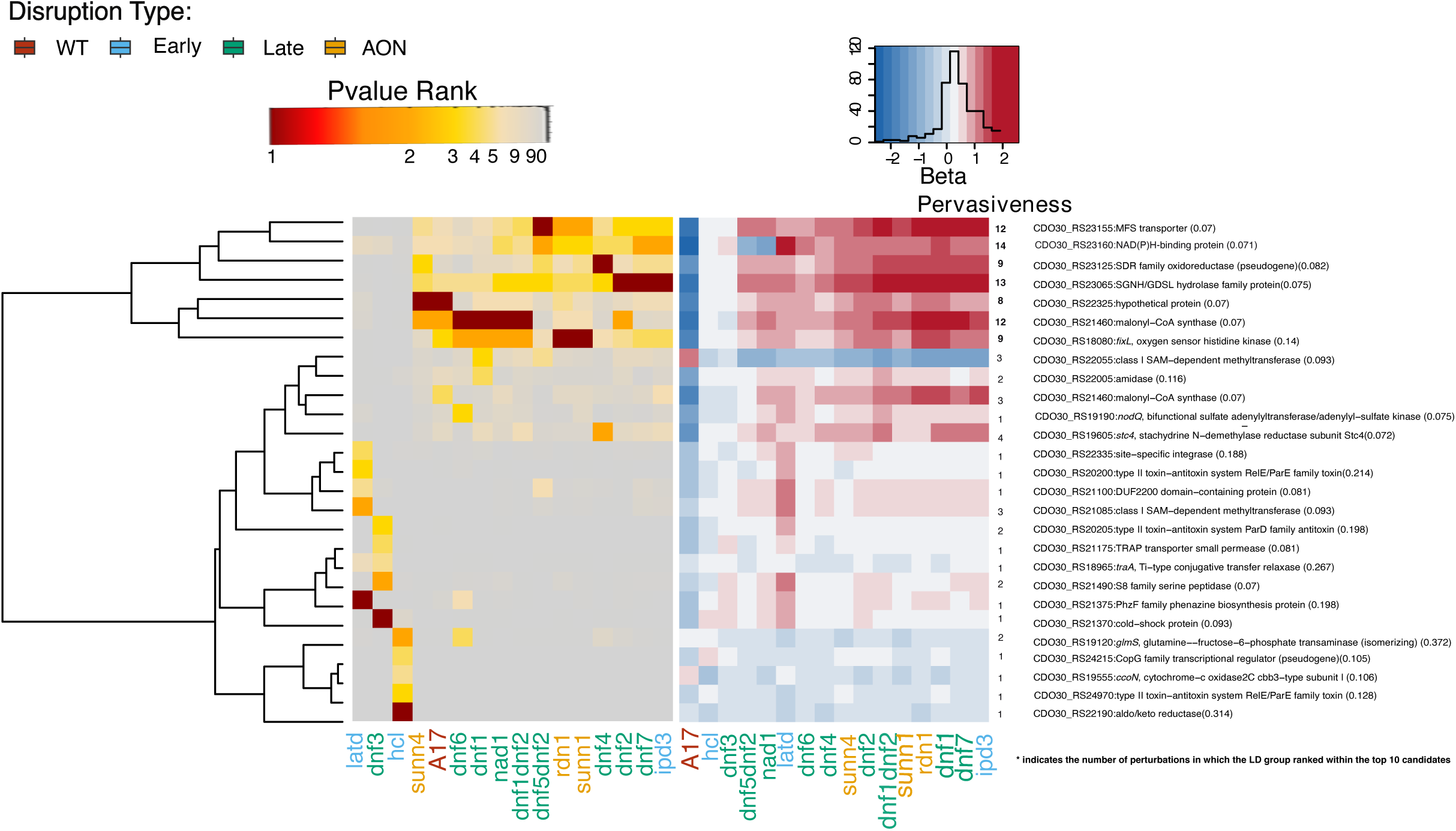
Rhizobial candidate genes underlying strain fitness shifts predominantly occur on pSymA. Heatmaps based on the inverse *P*-value ranks (red-yellow) and β values, which denote the effect sizes (red-blue) for the top five variant groups on the pSymA across the 15 host mutants and A17. Rows represent candidate variant groups, and columns represent host mutants. The host mutants are colour coded based on the stage of disruption they affect. Progressive dark shades (low *P*-value ranks) indicate a stronger association between the variant group and strain fitness shift in that host mutant. Heatmap based on the effect sizes (β values) for the same candidate variant groups was ordered to match the *p*-value ranking. Shades of red indicate host mutants where the corresponding variant has a positive effect on strain fitness shift, and shades of blue indicate negative effects as compared to A17. Colour intensity corresponds to effect magnitude. Alternate allele frequencies (i.e., not present in the reference genome USDA1106) for each candidate are also shown in parentheses next to the gene names. Pervasiveness of a variant group is defined as the number of times a variant group is among the top 10 candidates (based on the *P* value ranks) within a host mutant. A variant group is classified as “pervasive” if it is in the top 10 across six or more host genotypes. In contrast, a variant is classified as “limited effect” if it is in the top 10 candidates across five or fewer host genotypes. We have mentioned the number of times a variant appears in the top 10 under the “Pervasiveness” column in the figure and have indicated the pervasive genes in bold. (See Fig. S5 for the other replicons.)

However, the seven pervasive variants on pSymA were most interesting to us because they spanned similar hosts and had strong statistical support. These candidates included malonyl-CoA synthase, SGNH/GDSL hydrolase family protein, *fixL,* an oxygen sensor histidine kinase, a short-chain oxidoreductase, and a transport protein belonging to the MFS (Major Facilitator Superfamily). We validated one pervasive candidate, the malonyl-CoA synthase, by knocking it out in *S. meliloti* Rm1021 genetic background (See Supplemental Methods) and competing the mutant (RmND1439) against the wild type strain across four host genotypes (*sunn4*, *dnf3*, *dnf7*, and A17). RmND1439 was almost completely absent from nodules, indicating a functional version of this gene is crucial for strain fitness in all hosts tested (Fig. S7). This result supports the gene’s pervasive effect and confirms that our approach can identify rhizobial genes influencing strain performance. We further observed that the non-reference variant often increased strain fitness within the host mutants. In addition, we noted a trend where the pervasive variants that conferred higher relative fitness in the mutant hosts, tended to have lower fitness in wildtype A17. For example, the malonyl-COA synthase had an alternate allele associated with increased strain fitness (positive beta) in several host mutant backgrounds (*ipd3*, *dnf1*, *dnf2*, *dnf6*, *dnf7*, *nad1*, *rdn1*, and *sunn1*). In contrast, the same allele was associated with decreased fitness (negative beta) in the wild-type A17 (Fig. 4, Table S12). This pattern suggests a host-dependent allelic effect with the allele conferring a fitness advantage in mutant hosts becoming disadvantageous in the wild-type host.

Most limited-effect variant groups (82 of 94) were small (having fewer than three genes). The majority of these were on pSymB (Table S13). Those with strong statistical support (*P* value<10^-5^) included variants in or nearby genes coding two ABC transporters (*sunn1, ipd3, dnf1/dnf2, rdn1)*, glutamine synthetase (*latd)* methyltransferases (*nad1*), transcriptional regulators (*rdn1*), a couple of hypothetical proteins (*ipd3, dnf1/dnf2,dnf5/dnf2,dnf2,sunn1)*and genes involved in purine metabolism (*dnf3*). A rhizobial glutamine synthetase gene, which converts glutamate and ammonia into glutamine to assimilate nitrogen, was the strongest candidate for variation in strain fitness shift in the *latd* genotype, which encodes a nitrate transporter^59^. This was unexpected because rhizobial glutamine synthetases are generally thought to be strongly downregulated in nodules where host-driven nitrogen assimilation processes dominate^57^. To further investigate the potential functional relevance of these candidates, we utilized AlphaFold Multimer, coupled with Foldseek, to predict possible protein–NCR peptide interactions. Host NCR peptides are a logical starting point for putative plant protein-rhizobial protein interactions because 1) several of our hosts are NCR mutants^27^, 2) NCRs are known to have strain-specific effects^60^ and 3) the large NCR family is derived from antimicrobial peptides, but we only know what a handful of them do^61–63^. We found that more than half of the top rhizobial candidate genes associated with strain fitness shifts (20 out of 35) had a high likelihood of physically interacting with one or more NCR peptides (Table S13, S14). These included rhizobial candidates associated with fitness shifts in NCR-compromised mutants whose annotations suggest that rhizobia might adjust their membrane transport, redox balance, and surface defences when host NCR activity changes (e.g., xanthine dehydrogenase family protein subunit M, a Teicoplanin resistance protein-like, and an ABC transporter ATP-binding protein, which could be an NCR efflux protein according to Foldseek). These findings provide additional support for the biological relevance of these candidates in mediating rhizobium–host interactions.

### Correlations of host investment and benefit traits with rhizobial traits

Because mutualistic partners affect each other’s fitness, we were curious if any host investment or benefit traits were correlated with rhizobial fitness. At the level of individuals pots, nodule area, plant biomass, and root-to-shoot ratio covaried with nodule strain diversity and position on the first RDA axis describing shifts in strain relative fitness (Fig. 5A). Total nodulated area per plant covaried with the second RDA axis describing shifts in strain relative fitness. Perhaps unsurprisingly, the host trait that covaried with the number of CFUs released per plant was the number of nodules per plant. Across the first PC of trait space, we found the wild-type plants grouped with the AON mutants, *hcl*, and three *dnfs* (3,4,6). In contrast, the second PC tended to separate the early-acting, AON, and wildtype plants from the late-acting mutants. Genotypic trait correlations confirmed these patterns (Fig. 5B). Host mutants that were larger and had lower root-to-shoot ratios, (i.e., those that benefited more from rhizobial N-fixation), tended to release more rhizobia (CFUs) from individual nodules, more rhizobia on a whole-plant level, and had nodule strain communities that resembled those in A17 nodules (Fig. 5B). Most host traits correlated strongly with strain diversity and position along the first rhizobial relative fitness RDA axis; The exception was average nodule area/plant, which was most strongly correlated with host position on the second RDA axis. Details on mutation-specific effects on host traits can be found in Fig. S6 and Table S3. Our results suggest that symbiotic gene disruptions strongly shape both partners’ fitness in a manner that reinforces mutualism—mutations that reduce rhizobial fitness also reduce host fitness.

**Fig 5:**
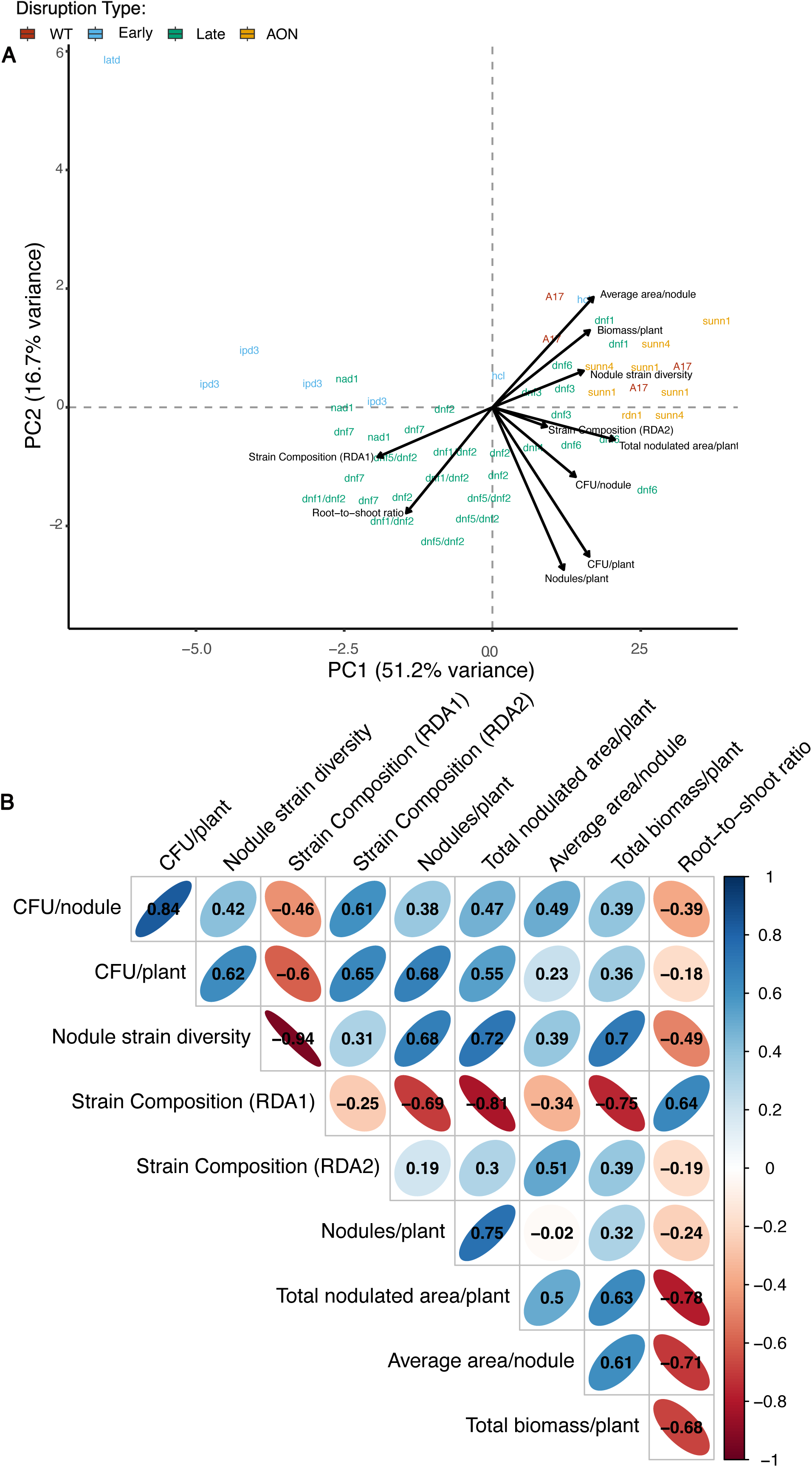
Covariation and correlation structure among host and rhizobial traits. (A) Principal component analysis (PCA) of host investment traits, benefit traits, and rhizobial fitness outcomes, conducted at the level of individual pots. Arrows represent the loadings of each trait on the first two principal components, which together explain 51.2% and 16.7% of the total variance, respectively. To avoid loss of representation due to missing data, values for two genotypes (*latd* and *rdn1*) were averaged across replicates prior to PCA, as these genes had incomplete trait data that would otherwise be excluded by listwise deletion. (B) All pairwise combinations of genotypic trait correlations (Pearsons). Each ellipse represents a pairwise Pearson correlation between traits, with ellipse elongation and color saturation indicating the strength and direction of the correlation (blue = positive, red = negative). Strong correlations may reflect shared genetic regulation or pleiotropy.

## Discussion

By combining a Select-and-Resequence (S&R) experiment with GWAS, we demonstrate that mutations in known host symbiotic genes create distinct environments that shape rhizobial strain success and identify rhizobial candidate genes underlying strain fitness shifts in response to host mutations. By applying a multi-strain inoculation approach here, we have created a more natural ecological setting to evaluate host-imposed selection on naturally occurring rhizobial genetic variants. Within this ecological framework, we found rhizobial genetic variants with both pervasive and limited effects across host mutants, and that symbiotic stage categories provide little clarity about how rhizobial genes influence strain fitness. We also found that mutations that increased rhizobial fitness tend to increase host investment and benefit traits, which highlight a directional effect where host genetic variations reshape selection on rhizobial partners. Our findings underscore the complexity of mutualism, where both host and rhizobial genomes jointly determine the outcomes of symbiosis at the community and individual strain levels, adding to knowledge of how *Medicago* hosts determine rhizobial fitness^16,30^.

### Host genetic mutations impact rhizobial absolute fitness

Legumes releasing rhizobia from their senescing nodules directly contribute to the soil rhizobial population^28,48,64^. We found that most mutations did not affect the number of rhizobia released from each nodule, and those that did, *ipd3* and *nad1*, released fewer rhizobia per nodule. The reduced number of rhizobia per nodule with these two mutants is not unexpected, given that disrupting *nad1* leads to overactive plant defence responses in nodules and disrupting *ipd3* traps rhizobia within infection threads, likely reducing survival^32,43^. Whereas our results suggest that the effects of host genetic changes on rhizobial fitness may sometimes align with known gene functions,, we also observed broader positive correlations between rhizobia released per nodule and average nodule area across mutations. Increasing the number of rhizobia the host supports represents one pathway to nurture robust populations of rhizobia, and identifying the host allelic variation that increases this amplification could set the stage for targeted interventions aimed at bolstering soil rhizobial populations in agricultural fields^65^. However, the host mutant lines used in this study were generated through forward genetic screens and, therefore, will likely carry additional background mutations. Thus, our aim is not to assign strict causality to individual mutations but rather to examine how alterations in the host genetic landscape broadly influence rhizobial fitness.

### Rhizobial fitness shifts depend on host genetics

A large group of NCR peptides is one of the major known host factors for altering strain survival and growth in nodules^63,66–68^. Although late-acting NCRs can be important determinants of host strain specificity in *M. truncatula*, our investigation reveals a more complex landscape^27,69^. Our analyses of two NCR mutations revealed differing effects on strain fitness within nodules. The *dnf7* mutant, which lacks functional NCR169^62^, was the only mutant background where we observedreduced nodule strain diversity, and it clustered with *ipd3,* which also supported smaller rhizobial population. In contrast, *dnf4*, which does not produce NCR211 and influences rhizobial differentiation during bacteroid formation^70^, showed shifts in strain relative fitness similar to *dnf3* and *dnf6,* suggesting these uncharacterized mutants may also involve NCR-related processes. A few rhizobia genes (*bacA, hrrP, yejABEF*) are known to affect the efficacy of NCRs^35,66,71^; however, none of those genes were top candidates in our GWAS, suggesting additional mechanisms of host restriction of rhizobia^61,72,73^.

### Rhizobial variants with pervasive and limited effects across host mutants

Among the ∼120 top rhizobial variants, ∼18% affected rhizobial fitness in six or more host mutant genotypes (pervasive genes), and ∼82% were associated with 5 or fewer host mutants(limited effect genes)., Pervasive effect alleles tended to exhibit fitness trade-offs; alleles linked to higher fitness in host mutants tended to be linked with lower fitness with wild-type hosts^16^. Many of the GWAS candidates were identified in only one or a few host mutant backgrounds, with the strongest associations on pSymB. Unlike the pervasive variants, most limited-effect variants had little to no effect on strain fitness in the wild-type background. Most candidate genes with pervasive associations were highly expressed in the nitrogen fixation zone (Zone III)^57^. For example, the *fixL* gene, which regulates the expression of nitrogen fixation genes in response to low oxygen levels, malonyl-CoA synthase that is induced in the rhizosphere of alfalfa, responsible for malonate catabolism within root nodules^77^, and a NAD(P)H-binding protein from the conserved NmrA domain family that shows high similarity to a transcriptional regulator, involved in nitrogen metabolite repression in fungi^78^. These genes represent strong candidates for uncovering novel mechanisms that influence symbiont performance across diverse host contexts^79^.

Beyond these candidates, most associations appeared host-specific, indicating that each mutated host background creates a unique challenge for rhizobia reproduction. Notable candidates include: A Class I SAM-dependent methyltransferase associated with strain fitness in *nad1* mutants, suggesting the importance of epigenetic modifications for rhizobial survival under host defences^32^, and *stc4*, encoding stachydrine N-demethylase reductase, which is linked to enhanced strain fitness in *dnf4* mutants, possibly by protecting differentiating rhizobia^70^.

Whereas these genes are candidates for altering strain fitness in some host mutant contexts and not others, our experiments do not pinpoint the stage at which each host gene exerts the strongest selective filter. One way to follow-up on these results is with assays comparing rhizobia mutated in the candidate genes identified in our GWAS on wildtype and mutant hosts, which would allow one to identify the selective stage—rhizosphere, infection thread, or nodule—where each gene exerts its influence on strain fitness.

### Considerations for interpreting mixed-strain experiments

Final strain fitness in nodules is a composite trait that reflects host genetics and microbial traits such as rhizosphere colonization, competition for infection, and proliferation within nodules. As such, shifts in relative strain fitness across host genotypes could also be influenced by differential competitive abilities among strains before infection of each host genotype. Thus, whereas our design allows us to detect which host symbiotic genes alter strain fitness, it does not allow us to disentangle the relative contributions of microbial competition versus host filtering. This limitation is especially relevant in complex communities, where strain-strain interactions influence host effects^21^. Whereas data suggest the effects of microbe × microbe interactions on strain relative fitness in our system are weak^26^, we caution that it is important to interpret our host-driven fitness shifts as emergent outcomes of host × microbe × microbe interactions. An additional consideration is that our experiment is designed to measure how growing plants of a single genotype affects the fitness of rhizobial strains. Linking these fine-scale host effects to long-term dynamics of rhizobial adaptation is an important overarching goal in plant-microbe interactions that requires incorporating additional ecological and evolutionary contexts, including the importance of soil selection^80^ or the cooccurrence of host genotypes across time and space^13,81^.

### Conclusion

Despite legumes being used for soil fertility improvement and actively managed for centuries, biological nitrogen fixation in agriculture remains inconsistent spatially and temporally^83–86^. This challenge has driven interest and investment in developing microbial inoculants to enhance crop yield, stress tolerance, and disease and pest management^20^. Rhizobia have been included in microbe-based bioinoculant formulations for centuries^79^. However, most of those strain selection decisions are dominated by strain performances observed under single-strain assays (inoculating a single plant with a single strain) in controlled environments. Thus, strains that show promise as potential bioinoculants under controlled conditions often fail to reach densities high enough to influence crop yields in realistic agricultural systems^87^. Formulating ways to sustain rhizobial fitness in agricultural fields is a critical avenue for plants to continue acquiring nitrogen from their associated rhizobia^88^.

Advances in host-mediated microbiome engineering have demonstrated that plant genes can be leveraged to assemble trait-optimized microbial communities, with the selection of beneficial microbes being dependent on the host genotype^85,89^. The findings in this study suggest a route towards expanding the search for host genes that alter rhizobial fitness. If natural allelic variants of these genes also favor colonization by high-performing symbionts, it will enable the development of “designer plants” with enriched mutualistic microbiomes^90^. Here, we identify host genes that drive strain-specific selection and show that even functionally similar mutations can have divergent effects on strain fitness in multi-strain contexts^21^. Whereas our study focused on loss-of-function mutations, studies on segregating natural genetic variation in *Medicago* suggest that specific host genes can differentially enrich rhizobia but find limited evidence for widespread balancing or positive selection^16^. Though we do not expect to find these loss-of-function mutations segregating in natural populations, this work lays a critical foundation: identifying the host genes that influence rhizobial fitness. A next step could be to systematically test the effect of naturally occurring alleles within these genes on rhizobial fitness, to pinpoint the variants that promote or reduce fitness and the mechanisms by which they do it.

## Acknowledgments

We thank Prof. Dong Wang of the University of Massachusetts Amherst for generously providing the *M. truncatula* mutant seeds. We also appreciate the assistance of Lillian Cherry and Jennifer E. Harris with the tedious task of picking and processing nodules. Prof. Katy Heath of the University of Illinois Urbana-Champaign and Dr. Mike Grillo (currently Assistant Professor at Loyola University, Chicago) for providing strains and sequencing data. This work was supported by the National Science Foundation grants IOS-1856744 to PT, NY, and LTB, and IOS-2243819, 1724993, 2243817, and 2243819 to PT and LTB. It was completed using computing resources provided by the Penn State Institute for Computational and Data Science, and sequencing was performed by the Huck Genomics Core Facility. LTB’s work was supported by the USDA National Institute of Food and Agriculture and Hatch Appropriations under Project #PEN04760 and Accession #1025611. The authors acknowledge the Huck Institutes’ Genomics Core Facility (RRID:SCR_023645) for providing library prep and sequencing services.

## Authorship Credits

LTB, NY, PT idea conception and funding, LTB experimental design, LTB and RGB plant care; RB, GF, LTB data collection for main experiment; GL, AG, BG constructed the malonyl-CoA synthase mutant, SG did the competition assays; VV and SS did the Alphafold Multimer and Foldseek analysis; SG, LTB, RGB, BE, JS, S&R and GWAS analysis; LTB, SG, RB drafted the manuscript; SG, LTB, RB, JS, BE, PT, NY, GF, BG, GL,VV, AG and SS for the manuscript revisions.

## Competing Interests

The authors declare no competing interests.

## Data Accessibility

All the raw reads from sequencing the rhizobial strains in the initial community and in the rhizobial genomic DNA extracted from the nodule pools of the individual host mutants have been deposited in NCBI under the Bioproject PRJNA401437 (Submission ID: SUB14622715). All raw data and analyses code have been uploaded to the Github (https://github.com/BurghardtLab/HostMutants2021_C86meliloti)

## Notes

https://github.com/BurghardtLab/HostMutants2021_C86meliloti

### Competing Interest Statement

The authors have declared no competing interest.

### Summary of Updates

1.Title Revision: Modified the manuscript title to remove causal phrasing and emphasize that host mutations cumulatively influence, rather than individual host genes determine, rhizobial fitness. 2.Additional GWAS analyses: Performed new GWAS analyses using Strain Fitness as the trait on individual host mutant lines and A17. All codes, figures, and tables are included in the GitHub repository and Supplement (Fig. S5; Tables S7,S8,S9,S10, S11). 3.Experimental validation of one GWAS candidate: Constructed the mutant of a single candidate gene and validated that this gene was needed for strain fitness. Added new experimental data validating candidate genes from GWAS and have included the data as a supplementary plot, with extended methodology, datasheets, and additional code uploaded to the GWAS repository. 4. Alphafold multimer analysis added (Tables S13,14) 5. Balanced and Precise Language: Clarified and toned-down language throughout the manuscript to describe host influence on strain fitness as the cumulative impact of all the host genetic perturbations rather than the effects of specific genes on strain fitness shifts.

