## Supplemental Methods and Figs for "Mutations in legume genes that influence symbiosis create a complex selective landscape for rhizobial symbionts"

Supplemental Tables

**Table S1-** Summary of *Medicago* mutant genotypes used

**Table S2** ANOVA results for the pairwise contrasts of response (traits) and the predictors.

**Table S3** Tukey HSD pairwise comparisons of A17 (wild-type) and each host mutant and the corresponding *P* values. (provided as `.csv` file)

**Table S4-** RDA paired with PERMANOVA analysis of nodule isolate fitness across the disruption types (Early, Late, and AON).

**Table S5-** RDA paired with PERMANOVA analysis of nodule isolate fitness across the genotypes

**Table S6-** List of the top five candidate genes across the different hosts, underlying strain fitness shift across chromosome, pSymA, pSymB (*P* values are indicated in the p_lrt column) (provided as `.tsv` file)

**Table S7-** List of the top five candidate genes across the different hosts, underlying strain fitness across chromosome, pSymA, pSymB (*P* values are indicated in the p_lrt column) (provided as `.tsv` file)

**Table S8**- List of the top 5 common genes underlying strain fitness across A17 and the host mutants (provided as `.tsv` file)

**Table S9**- List of the top 5 unique genes underlying strain fitness in the host mutants (provided as `.tsv` file)

**Table S10**- List of the top 5 common genes underlying strain fitness and strain fitness shifts across the host mutants (provided as `.tsv` file)

**Table S11**- List of the top five unique candidate genes underlying strain fitness shift in the host mutants (provided as `.tsv` file)

**Table S12-** Cross ranking of top five candidates underlying strain fitness shift in each host mutant (provided as `.tsv` file)

**Table S13-** AlphaFold Multimer predicted NCR targets of the rhizobial candidate genes (provided as .xlsx file)

**Table -S14**: Key values (pae ,ptm and pITM) from the Alpha Fold analysis of GWAS candidates (provided as .xlsx file)

**Table S15-** Strains used for functional validation of the GWAS candidate Malonyl CoA synthase.

**Table S16**- Plasmids and primers used for functional validation of the GWAS candidate malonyl CoA synthase.

<u>Supplemental Methods</u>

***Generating the malonyl-CoA synthase mutant in S.meliloti Rm1021:***

*Strain Growth Conditions:* Bacterial strains and plasmids used in this work are listed in Table S15. *Escherichia coli* strains were grown at 37°C in Luria-Bertani (LB) medium supplemented with either chloramphenicol (Cm) at 5 µg $mL^{-1}$ or gentamicin (Gm) at 20 µg $mL^{-1,}$ as appropriate. A concentration of 40 µg $mL^{-1}$ of 5-bromo-4-chloro-3-indolyl-beta-D-galactopyranoside (X-gal) was used when blue-and-white screening via *lacZ* was implemented. *S.meliloti* strains were grown at 28°C in LB medium supplemented with 2.5 mM $MgCl_2$ and 2.5 mM $CaCl_2$ (LBMC). When appropriate, concentrations of streptomycin (Sm) 200 µg $mL^{-1}$ and gentamicin (Gm) 60 µg $mL^{-1}$ were used when growing *S.meliloti.*
*Creation of malonyl-CoA synthase mutant:* Primers used for the creation of pNDMS802 and RmND1439 are listed in Table S16. To construct pNDMS802 an amplified internal fragment of the malonyl-CoA synthase gene was integrated into pNDGG014 via Golden Gate cloning [1]. The construct was transformed via heat-shock into DH5-Alpha and screened via antibiotics, blue-and-white screening, and PCR. The verified plasmid was then introduced into Rm1021 by conjugation via triparental mating between the helper plasmid MT616, donor (pNDMS802), and recipient[2]. Integration of pNDMS802 into the Rm1021 genome was verified via PCR and amplification across the genome and plasmid junctions using primers listed in Table S16.
*Competition assay:* Rm1021 and RmND1439 were grown individually in liquid LBMC media for 3.5 days in a 28 °C shaker. Both strains were combined in equal proportions to create the initial community, which was used to inoculate the host genotypes. To determine strain representation in the initial community, we plated four replicate serial dilutions of the strain

mixture on LBMC and LBMC+gentamicin plates and calculated the proportion of the mutants in the initial culture by comparing the colonies growing in the presence and absence of antibiotics. Plant germination and plant care were performed as described earlier. We inoculated 16 replicates per host and had uninoculated control plants from each host category to monitor cross-contamination. Following 8 weeks post-inoculation, the plants were harvested, the nodules collected, photographed, homogenised and serially diluted. We plated equal volumes of the serial dilutions from each of the host plants on LBMC plates with and without antibiotics to count the proportion of Rm ND1439 in the nodules. Both the wildtype (Rm1021) and the mutant (RmND1439) should grow on the LBMC plates, but only the mutant strain (RmND1439) will survive on the LBMC+gentamicin. Rm ND1439 fold change for each dilution replicate was calculated as $\log_2$(Colonies on LBMC+gentamicin plates /Colonies on LBMC).

***Putative annotation of rhizobial candidate genes using Foldseek:***

The rhizobial candidate protein sequences were matched to their entries in the AlphaFold Database [3,4] , which were then loaded onto the FoldSeek Search web server [5]. Hits with RMSD values less than 3 Å and sequence alignment scores more than 50% were considered significant for the purpose of functional annotation. The most structurally similar and sequence-aligned hits were selected to be the functional equivalent of the query protein.

***Prediction of protein-protein interactions of the candidate genes with NCR peptides using AlphaFold-Multimer***

Protein complexes were modelled using AlphaFold-Multimer v3 [3,6]). Interactions were modelled between the query protein of interest (rhizobial candidate gene) against all NCR peptides as the bait set of sequences. Query and bait sequences were arranged in FASTA format in separate files. ColabFold was run on all monomer sequences first, and sequences with a pLDDT score below 70% were excluded [7]. Multiple sequence alignments (MSAs) were generated for each of the individual queries and baits. Query and bait MSAs were combined to generate multimer MSAs for every possible pair of query and bait combinations. Then ColabFold was run on each of the multimer MSAs to generate plots of (coverage, pLDDT, pAE), a structure file (.pdb), and a scores file (.json) with key metrics of the interaction like plddt, pTM, ipTM, and pAE. The pAE_interaction score for a given prediction is calculated by taking the average of the off-diagonal blocks from the pAE matrix generated. Pae_interaction is the average predicted alignment error of interchain residue pairs. PAE is a measure of how confident AlphaFold2 is in the relative position of two residues within the predicted structure. Interchain residue pairs are pairs of amino acid residues, one from each of two separate protein chains, that are physically close to each other in the 3D structure of a protein complex. The lower this score is, the better, and a PAE<10 is considered significant. The pTM (predicted Template Modeling) score is a metric (ranging 0-1) that assesses the overall accuracy of the predicted protein complex structure, with higher scores indicating better quality. A pTM score above 0.5 suggests that the overall fold of the complex is likely similar to the true structure. iptm: (interface pTM) specifically measures the accuracy of the predicted relative positions of the protein subunits at the binding interface. pAE matrix generated.

***Nodule-enrichment scores***

Nodule enrichment scores for each of the bacterial query proteins were obtained by entering the gene identifiers (e.g., Sma0001) into the Symbimics database (https://iant.toulouse.inra.fr/symbimics/), which hosts an integrated analysis of bacterial gene expression in symbiotic root nodules using laser-capture microdissection coupled to RNA sequencing [8]. The nodule enrichment score is defined as a measure of deseq-normalized RNA-seq reads from the whole-organ analysis experiment with rRNA-depleted RNA.

**Table S1**- Summary of *Medicago* mutants used. All mutants were in the A17 background.

| Mutant line | Symbiosis stage affected | Host investment /benefit[1] | Reference |
|---|---|---|---|
| *hcl* | Early | +/- | [1] |
| *ipd3* | Early | +/- | [9,10] |
| *latd* | Early | +/- | [11] |
| *dmi1-2* | Early | -/- | [12] |
| *dmi2-3* | Early | -/- | [12] |
| *dmi3-1* | Early | -/- | [13] |
| *dnf1* | Late | +/- | [14] |
| *dnf2* | Late | +/- | [15] |
| *dnf3* | Late | +/- | [16] |
| *dnf4* | Late | +/- | [16] |
| *dnf5* | Late | +/- | [16] |
| *dnf6* | Late | +/- | [16] |
| *dnf7* | Late | +/- | [16] |
| *dnf1/dnf2* | Late | +/- | Unpublished (Dong Wang) |
| *dnf5/dnf2* | Late | +/- | Unpublished (Dong Wang) |
| *nad1-3* | Late | +/- | [17] |
| *sunn1* | AON | +/- | [18] |
| *sunn4* | AON | +/- | [18] |
| *rdn1* | AON | +/- | [19,20] |

[1]Host investment/benefit (+/-) means positive nodulation(+) and no biomass increase (-). The information was compiled from single-strain data obtained from previous publications

**Table S2**- ANOVA results for the pairwise contrasts of response (traits) and the predictors.

| Response | Pred. [1,*a] | %Exp [2] | Df [3] | Fval [3] | Pval [4] | $R_{adj}$ [5] | Pred | %Exp | Df | Fval | Pval | $R_{adj}$ |
|---|---|---|---|---|---|---|---|---|---|---|---|---|
| Nodules per plant | Geno. | 66.3 | 15 | 7.04 | 0 | 0.59 | Type | 42.29 | 3 | 14.96 | 0 | 0.39 |
| Nodules per plant | Rep. | 4.8 | 4 | 1.92 | 0.12 | NA | Rep. | 3.06 | 4 | 0.81 | 0.52 | NA |
| Nodules per plant | Res. | 28.8 | 46 | NA | NA | NA | Res. | 54.65 | 58 | NA | NA | NA |
| Average area per nodule | Geno. | 61.1 | 15 | 4.91 | 0 | 0.51 | Type | 24.53 | 3 | 6.37 | 0.001 | 0.24 |
| Average area per nodule | Rep. | 5.6 | 4 | 1.70 | 0.16 | NA | Rep. | 8.71 | 4 | 1.70 | 0.16 | NA |
| Average area per nodule | Res. | 33.2 | 40 | NA | NA | NA | Res. | 66.75 | 52 | NA | NA | NA |
| Total nodulated area per plant | Geno. | 84.6 | 15 | 22.56 | 0 | 0.85 | Type | 60.86 | 3 | 32.78 | 0 | 0.63 |
| Total nodulated area per plant | Rep. | 5.3 | 4 | 5.33 | 0 | NA | Rep. | 6.96 | 4 | 2.81 | 0.03 | NA |
| Total nodulated area per plant | Res. | 10 | 40 | NA | NA | NA | Res. | 32.18 | 52 | NA | NA | NA |
| Total biomass per plant | Geno. | 80 | 18 | 17.65 | 0 | 0.79 | Type | 24.92 | 3 | 9.03 | 0 | 0.22 |
| Total biomass per plant | Rep. | 4.2 | 5 | 3.39 | 0 | NA | Rep. | 4.25 | 5 | 0.92 | 0.47 | NA |
| Total biomass per plant | Res. | 15.6 | 62 | NA | NA | NA | Res. | 70.83 | 77 | NA | NA | NA |
| Root-to-shoot ratio | Geno. | 89.1 | 18 | 36.34 | 0 | 0.88 | Type | 39.28 | 3 | 17.28 | 0 | 0.36 |
| Root-to-shoot ratio | Rep. | 2.3 | 5 | 3.48 | 0 | NA | Rep. | 2.38 | 5 | 0.63 | 0.679 | NA |
| Root-to-shoot ratio | Res. | 8.4 | 62 | NA | NA | NA | Res. | 58.34 | 77 | NA | NA | NA |
| CFU per nodule | Geno. | 60.5 | 15 | 5.43 | 0 | 0.54 | Type | 13.24 | 3 | 3.19 | 0.03 | 0.14 |
| CFU per nodule | Rep. | 7.4 | 4 | 2.51 | 0.05 | NA | Rep. | 10.76 | 4 | 1.95 | 0.116 | NA |
| CFU per nodule | Res. | 31.9 | 43 | NA | NA | NA | Res. | 76.00 | 55 | NA | NA | NA |
| CFU per plant | Geno. | 83.6 | 15 | 22.30 | 0 | 0.84 | Type | 40.37 | 3 | 14.90 | 0. | 0.42 |
| CFU per plant | Rep. | 5.0 | 4 | 5.09 | 0 | NA | Rep. | 8.17 | 4 | 2.26 | 0.07 | NA |
| CFU per plant | Res. | 11.2 | 45 | NA | NA | NA | Res. | 51.46 | 57 | NA | NA | NA |
| Nodule isolate diversity | Geno. | 58.3 | 15 | 3.99 | 0 | 0.44 | Type | 22.78 | 3 | 5.37 | 0 | 0.19 |
| Nodule isolate diversity | Rep. | 4.6 | 4 | 1.19 | 0.33 | NA | Rep. | 6.52 | 4 | 1.15 | 0.34 | NA |
| Nodule isolate diversity | Res. | 37 | 38 | NA | NA | NA | Res. | 70.70 | 50 | NA | NA | NA |
| Nodule isolate fitness | Geno. | 41.4 | 15 | 2.03 | 0 | 0.23 | Type | 12.78 | 3 | 2.67 | 0 | 0.09 |
| Nodule isolate fitness | Rep. | 6.8 | 4 | 1.26 | 0.05 | NA | Rep. | 7.36 | 4 | 1.15 | 0.1 | NA |
| Nodule isolate fitness | Res. | 51.6 | 38 | NA | NA | NA | Res. | 79.86 | 50 | NA | NA | NA |

[1]Pred.= Predictors.
[2]%Exp = percentage of total variance explained by each predictor.
[3]Df = degrees of freedom.
[4]Fval = F statistic from the ANOVA model.
[5]Pval = significance level of the predictor effect.
[6]Radj = adjusted $R^2$ for the full model.
[*a] Geno=Genotype; Rep.=Replicate; Res.=Residuals
*NA = not applicable.

**Table S4-** RDA paired with PERMANOVA analysis of nodule isolate fitness across the disruption types (Early, Late, and AON).

| **Trait** | **Predictor** | **R.adj**[1] | **Prop.Var**[2] | **F.val**[3] | **P.val**[4] | **Class** |
|---|---|---|---|---|---|---|
| Nodule_isolate_fitness | AON | 0.04 | 0.1137 | 1.66 | 0.02 | Change |
| Nodule_isolate_fitness | Early | 0.05 | 0.1356 | 1.72 | 0.03 | Change |
| Nodule_isolate_fitness | Late | 0.03 | 0.0619 | 2.37 | 0 | Change |

[1]Radj = adjusted $R^2$ for the full model.
[2]Prop.Var = Proportion of total variance explained by each predictor.
[3]Fval = F statistic from the ANOVA model.
[4]Pval = significance level of the predictor effect.

**Table S5**- RDA paired with PERMANOVA analysis of nodule isolate fitness across the genotypes

| **Trait** | **Predictor** | **R.adj[1]** | **Prop.Var[2]** | **F.val[3]** | **P.val[4]** | **Class** |
|---|---|---|---|---|---|---|
| Nodule_isolate_fitness | *hcl* | 0.0482 | 0.18 | 1.35 | 0.10 | None |
| Nodule_isolate_fitness | *ipd3* | 0.1519 | 0.27 | 2.25 | 0.03 | Change |
| Nodule_isolate_fitness | *latd* | 0.3102 | 0.48 | 2.79 | 0.2 | None |
| Nodule_isolate_fitness | *nad1* | 0.1467 | 0.26 | 2.20 | 0.03 | Change |
| Nodule_isolate_fitness | *dnf1* | 0.0331 | 0.17 | 1.23 | 0.21 | None |
| Nodule_isolate_fitness | *dnf2* | 0.141 | 0.26 | 2.14 | 0.02 | Change |
| Nodule_isolate_fitness | *dnf3* | 0.0894 | 0.21 | 1.68 | 0.02 | Change |
| Nodule_isolate_fitness | *dnf4* | 0.0995 | 0.27 | 1.55 | 0.06 | None |
| Nodule_isolate_fitness | *dnf6* | 0.1043 | 0.23 | 1.81 | 0.03 | Change |
| Nodule_isolate_fitness | *dnf7* | 0.1589 | 0.27 | 2.3 | 0.02 | Change |
| Nodule_isolate_fitness | *dnf1/dnf2* | 0.1934 | 0.30 | 2.67 | 0.02 | Change |
| Nodule_isolate_fitness | *dnf5/dnf2* | 0.1341 | 0.25 | 2.08 | 0.03 | Change |
| Nodule_isolate_fitness | *rdn1* | 0.0473 | 0.20 | 1.29 | 0.14 | None |
| Nodule_isolate_fitness | *sunn1* | 0.0493 | 0.18 | 1.36 | 0.08 | None |
| Nodule_isolate_fitness | *sunn4* | 0.094 | 0.22 | 1.72 | 0.06 | None |

[1]Radj = adjusted $R^2$ for the full model.
[2]Prop.Var = Proportion of total variance explained by each predictor.
[3]Fval = F statistic from the ANOVA model.
[4]Pval = significance level of the predictor effect

**Table S15**- Strains used for functional validation of the GWAS candidate Malonyl CoA Synthase.

| **Strain** | **Description** | **Reference** |
|---|---|---|
| ***Sinorhizobium meliloti*** | | |
| **Rm1021** | SU47 *str*-21; $Sm^R$ | [21] |
| **RmP110** | Wildtype Derived from Rm1021 with Corrected *pstC* Allele; $Sm^R$ | [22] |
| **RmND1439** | RmP110 with pNDMS802 Integrated (*malCoA*::*Gm)*; $Sm^R$ $Gm^R$ | This Work |
| ***Escherichia coli*** | | |
| **DH5-Alpha** | *fhuA2Δ(argF-lacZ)U169 phoA glnV44 Φ80Δ(lacZ)M15 gyrA96 recA1 relA1 endA1 thi-1 hsdR17* | **New England BioLabs** |
| **MT616** | MM294A *recA-56* (pRK600), Mobilizer; $Cm^R$ | [2] |
| **Plasmid** | **Description** | **Reference** |
| **pNDGG014** | BEVA (Gm and No FRT) P1 (pOGG004_noT) P2 (pOGG009) P3 (pTH3307 (pMBI via BsmBI)) P4 (pOGG013); $Gm^R$ | [1] |
| **pNDMS802** | pNDGG014 with Rm1021pSymA (GenBank: AE006469.1) nt 82,845 – 83,381 (7F-Mal/7R-Mal) via BsaI Golden Gate Cloning; $Gm^R$ | This Work |

**Table S16:** Primers used for functional validation of the GWAS candidate Malonyl CoA Synthase

| **Primers Used for Construction of pNDMS802** | | | | |
|---|---|---|---|---|
| **Primer** | **Forward(5'-3')** | **Primer** | **Reverse (3'-5')** | **Location** |
| **7F-Mal** | TTTGGTCTCAGGAGCTTCACCTCGAT CATCCCCG | **7R-Mal** | TTTGGTCTCTAGCGCTGACGCAT GACAACCTCCT | Rm1021 pSymA nt 82,845 – 83,381 |

| **Primers Used to Verify the Integration of pNDMS802into the Genome** | | | |
|---|---|---|---|
| **Primer** | **Forward** | **Primer** | **Reverse** |
| **7-V-F (Genome Based Forward Verification)** | CCAATCACAGCCGTTTCCAC | **GLND218 (Plasmid Based Reverse Verification)** | TGTAGGTATCTCAGTTCGGT |
| **GLND217 (Plasmid Based Forward Verification)** | GCTGCTCCATAACATCAAAC | **7-V-R (Genome Based Reverse Verification)** | CGAAGGAGGGGATTGCGAAG |

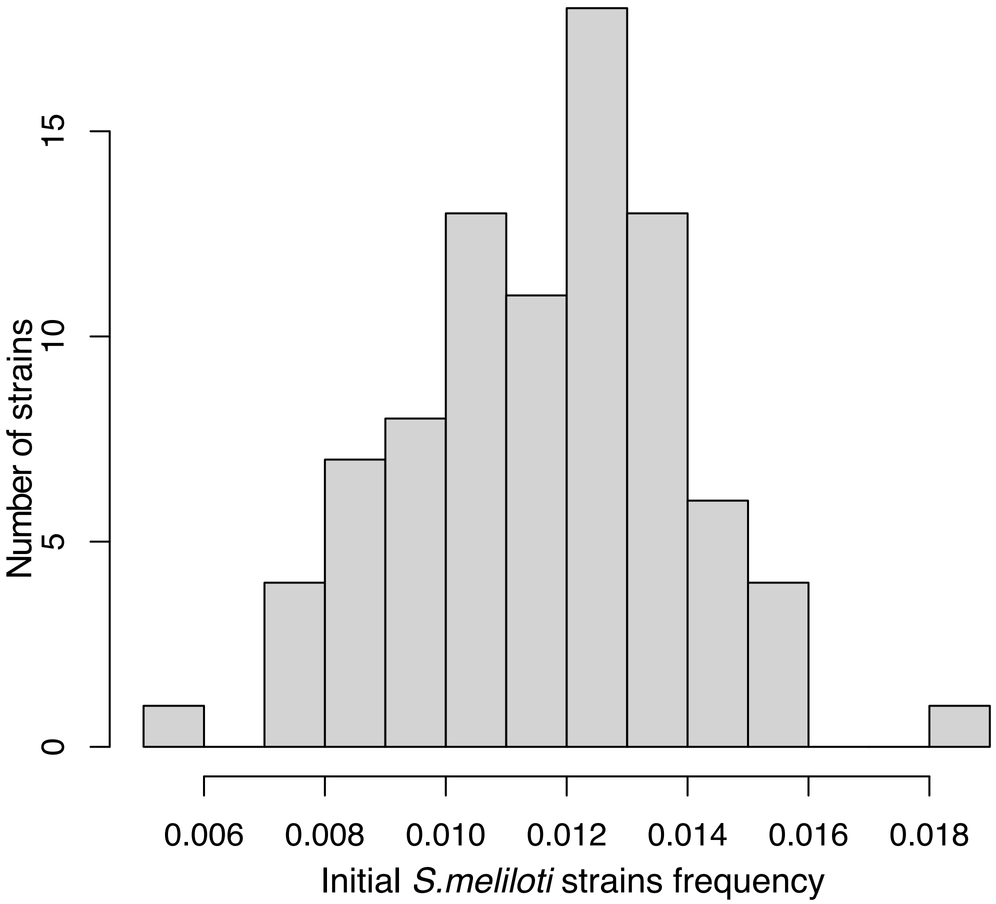

**Fig S1**: Histogram showing the distribution of the frequencies of 86 *S. meliloti* strains in the initial community used to inoculate the A17 wild type and the 18 mutant hosts.

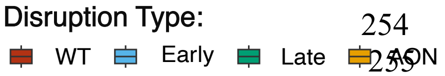

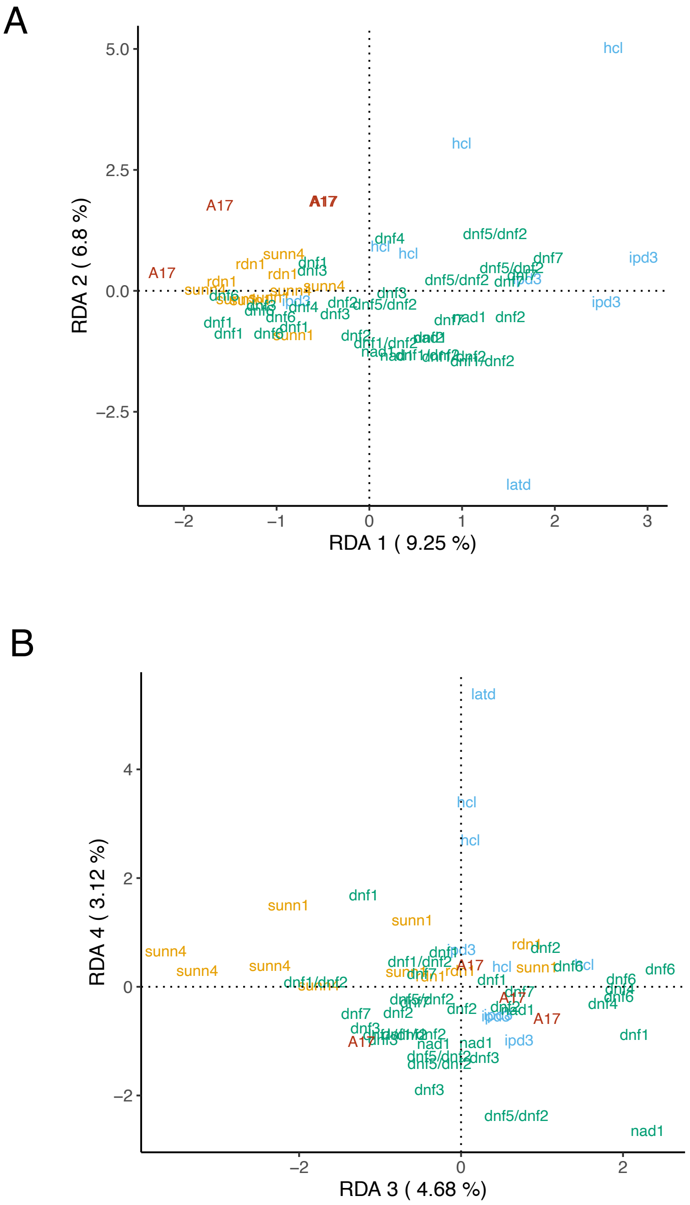

**Fig S2:** Host genotype explains 36% of the total variation in strain relative fitness. Redundancy Analysis (relative strain fitness ~ host genotype) showing the first and second RDA axis (A) and the third and fourth RDA axis (B). Points denote replicate pots, colours are coded based on the symbiosis category, and individual genotypes are labelled.

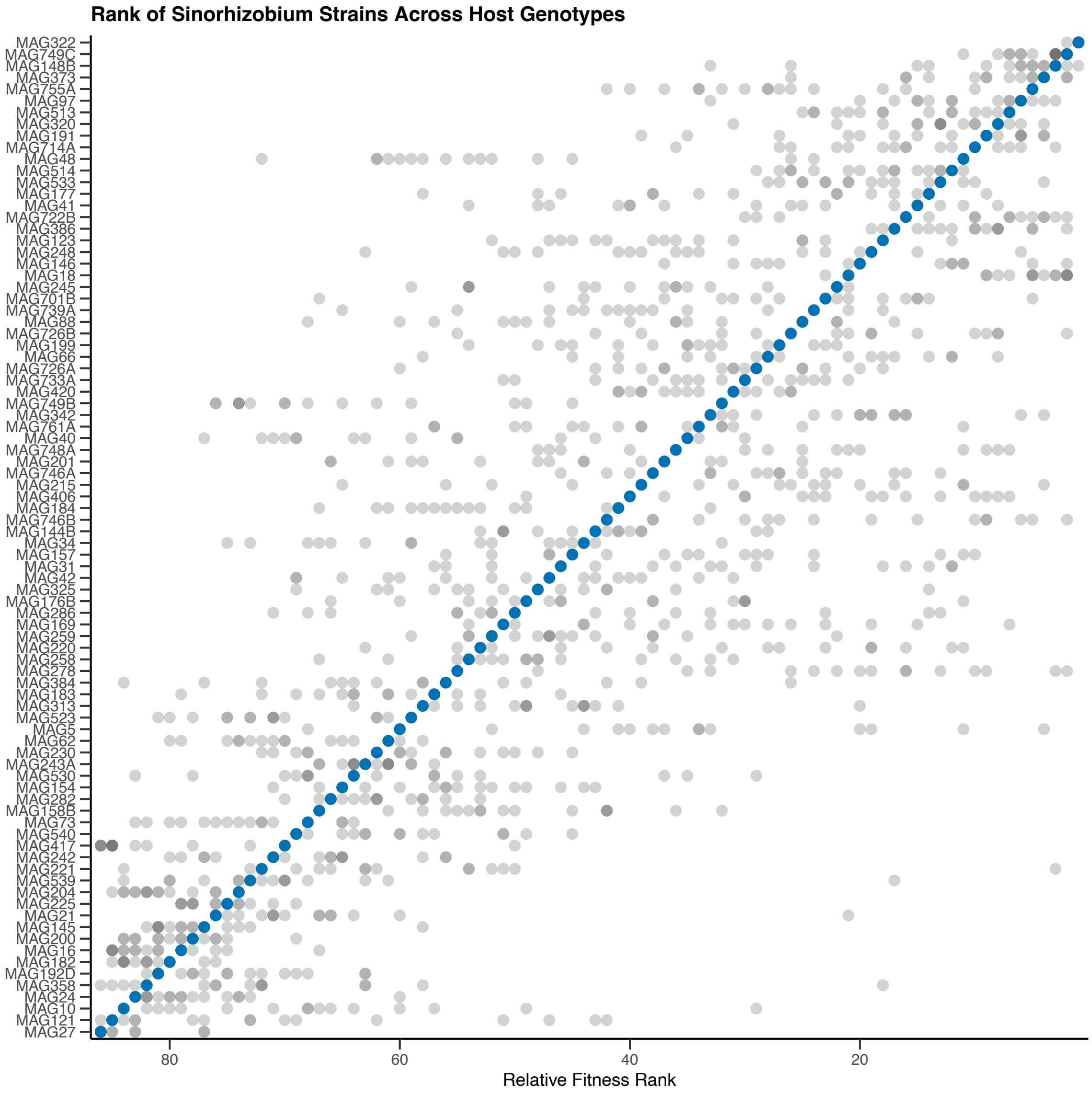

**Fig S3:** Strain fitness ranks compared across host genotypes in a scatterplot where each point shows the rank of a strain's fitness within a host genotype. Smaller numbers along the x-axis indicate a higher fitness rank, with lower ranks indicating higher fitness. Strains (y-axis) are ordered based on their rank in the A17 genotype (blue points). Gray points represent the ranks of each strain across all host genotypes. The x-axis is reversed so that higher fitness ranks (lower numbers) appear on the left.

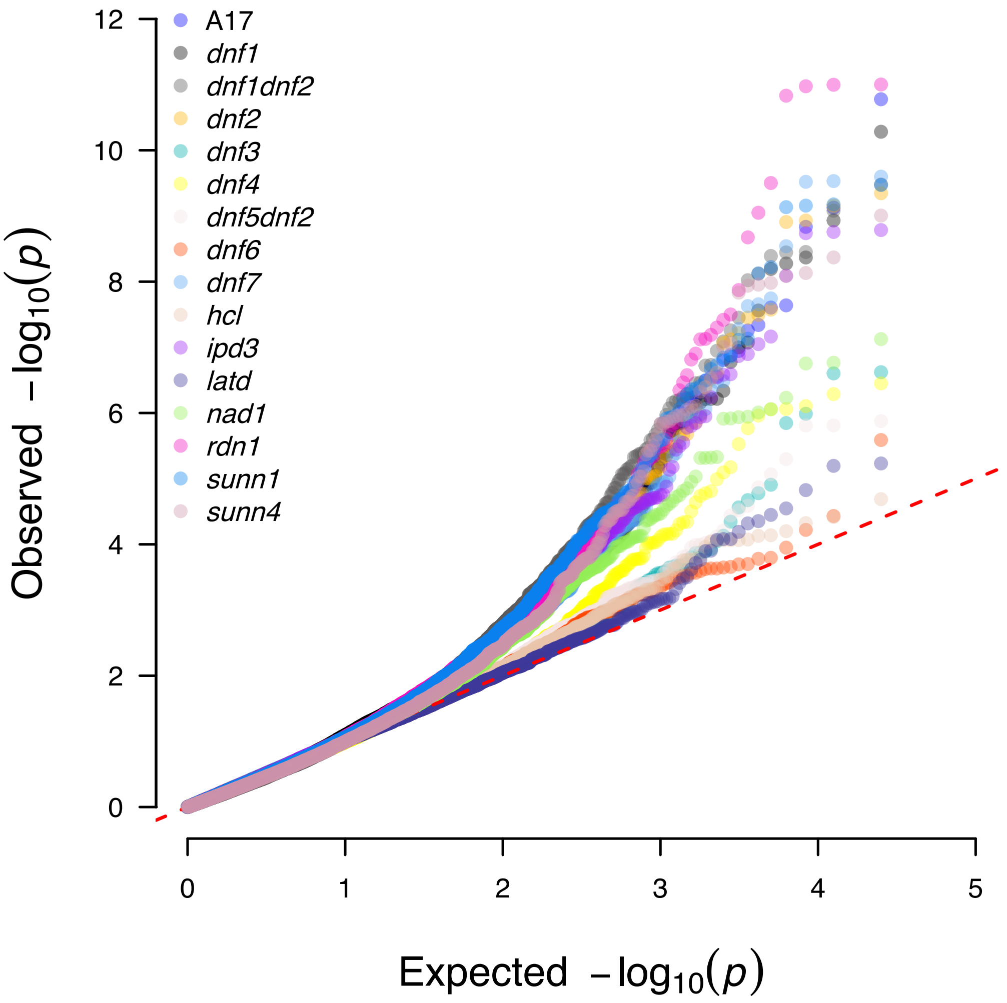

**Fig S4:** Quantile-quantile plot corresponding to the Manhattan plot in Fig. 3B, showing the empirically observed quantiles (y-axis) vs. the expected quantiles (x-axis) of the effects of the identified candidate genes on strain fitness shifts of the 86 *S.meliloti* strains in the different host mutants (colour coded as per the key listed). The null distribution is shown as a dotted red line.

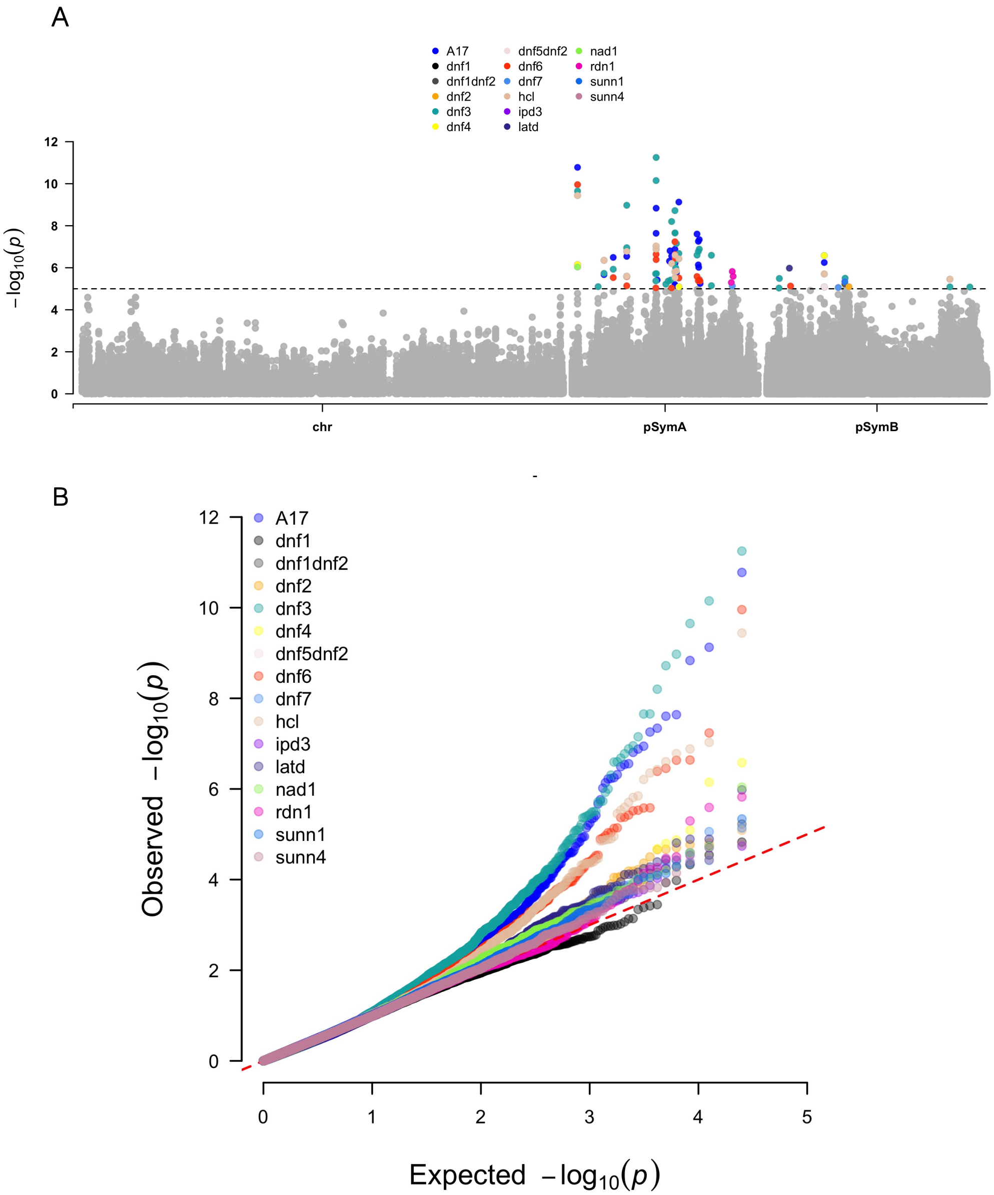

**Fig S5:** **(A)** Manhattan plot indicating the SNP (single-nucleotide polymorphism) positions, based on S. *meliloti* USDA1106 association with the relative strain fitness across the 15 host mutants and in A17. The dotted line indicates the $p$ value threshold used for identifying candidate genes ($p < 10^{-5}$), and significant associations are color-coded according to host mutation and A17. The GWAS was run separately on each replicon to account for different population structures. (Refer Table S7 for further details). **(B)** Quantile-quantile plot corresponding to the Manhattan plot in (A), showing the empirically observed quantiles (y-axis) vs. the expected quantiles (x-axis) of the effects of the identified candidate genes on strain fitness of the 86 *S.meliloti* strains in the different host mutants (colour coded as per the key listed). The null distribution is shown as a dotted red line.

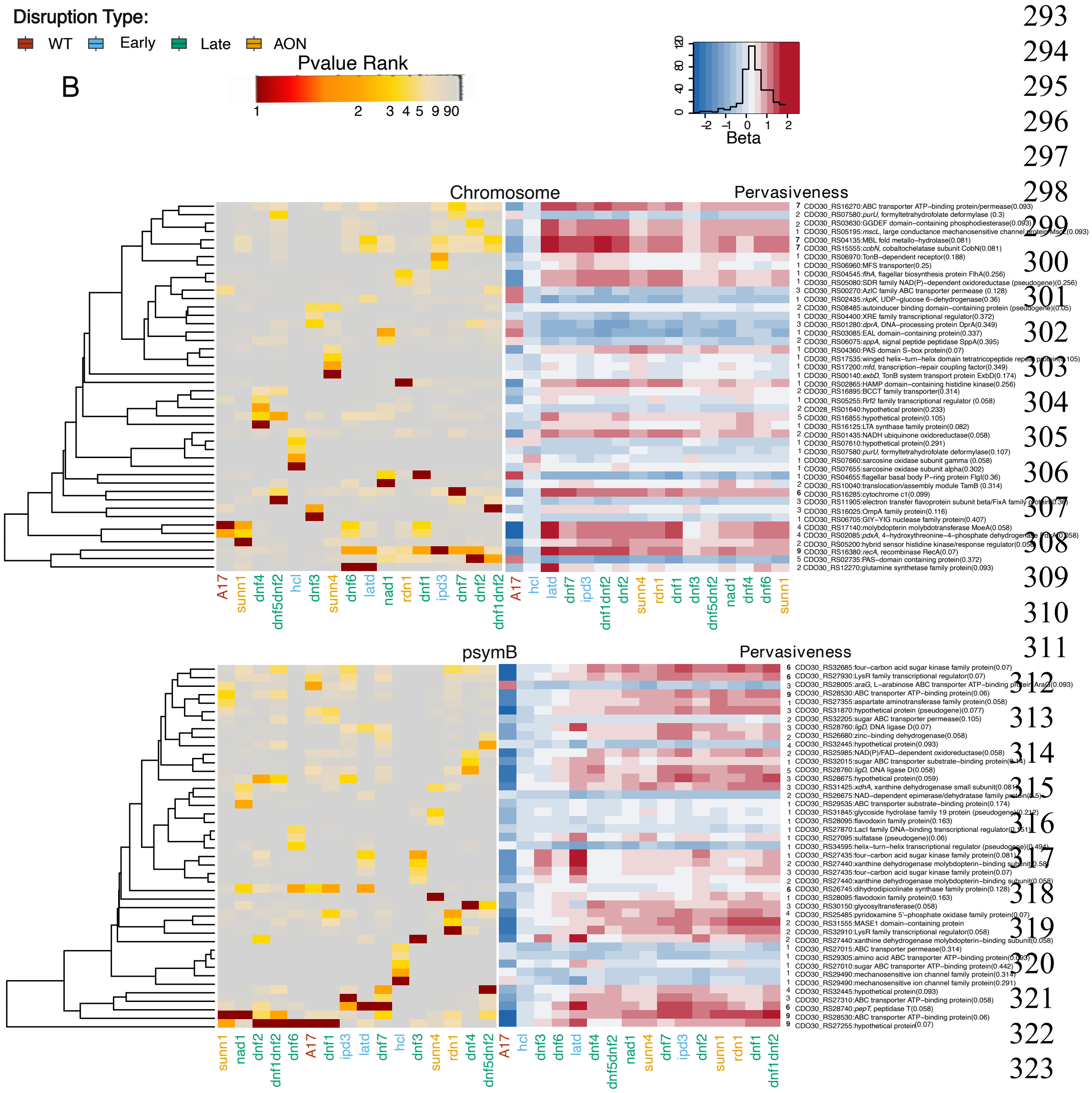

**Fig S6:** Heatmaps based on the inverse *P value* ranks (red-yellow) and effect sizes (red-blue) for the top five variant groups on the pSymB (A) and Chromosome (B) across the 15 host mutants and A17. Rows represent candidate variant groups, and columns represent host mutants. Progressive dark shades (low *P value* ranks) indicate a stronger association between the candidate and strain fitness shift in that host mutant. Heatmap based on the effect sizes (β values) for the same candidate variant groups was ordered to match the *P value* ranking. Shades of red indicate host mutants where the corresponding variant had a positive effect on strain fitness shift, and shades of blue indicate negative effects as compared to A17. Colour intensity corresponds to effect magnitude. The host mutants are colour coded based on the stage of symbiosis they disrupt. Alternate allele frequencies (i.e., not present in the reference genome USDA1106) for each candidate are also shown in parentheses next to the gene names. Pervasiveness of a variant group is defined as the number of times a variant group is among the top 10 candidates (based on the *P value* ranks) within a host mutant. A variant group is classified as "pervasive" if it is in the top 10 across six or more host genotypes, while a variant is classified as "limited effect" if it is in the top 10 candidates across five or fewer host genotypes. We have mentioned the number of times a variant appears in the top 10 under the "Pervasiveness" column in the figure and have indicated the pervasive genes in bold (Refer to Table S6 for further details)

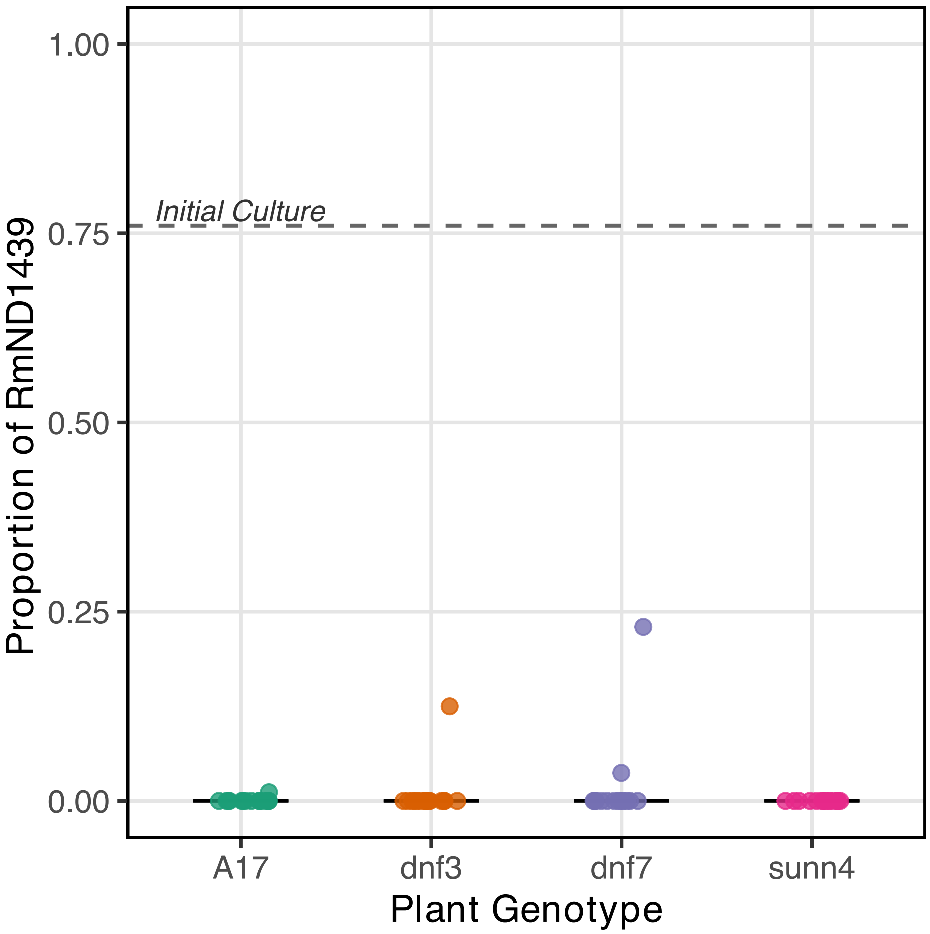

**Fig S7:** Proportion of RmND1439 (vs. *R*m1021) inside nodules of A17 and three host mutant genotypes-*dnf3, dnf7* and *sunn4*. Each point represent CFU counts obtained from ~14 nodules (A17), ~8.5 nodules(*dnf3*), ~17 nodules(*dnf7*), ~42 nodules (*sunn4*) from each of 15 replicate plants. The dashed line indicates the frequency of the mutant strain in the initial mixture (In this case, 75% of the initial mixture was RmND1439). Note that the y-axis is plotted on the $\log_2$ scale but labelled with proportions. A value of 1 indicates all colonies are RmND1439 mutants, whereas a value of 0 indicates that all colonies are Rm1021.

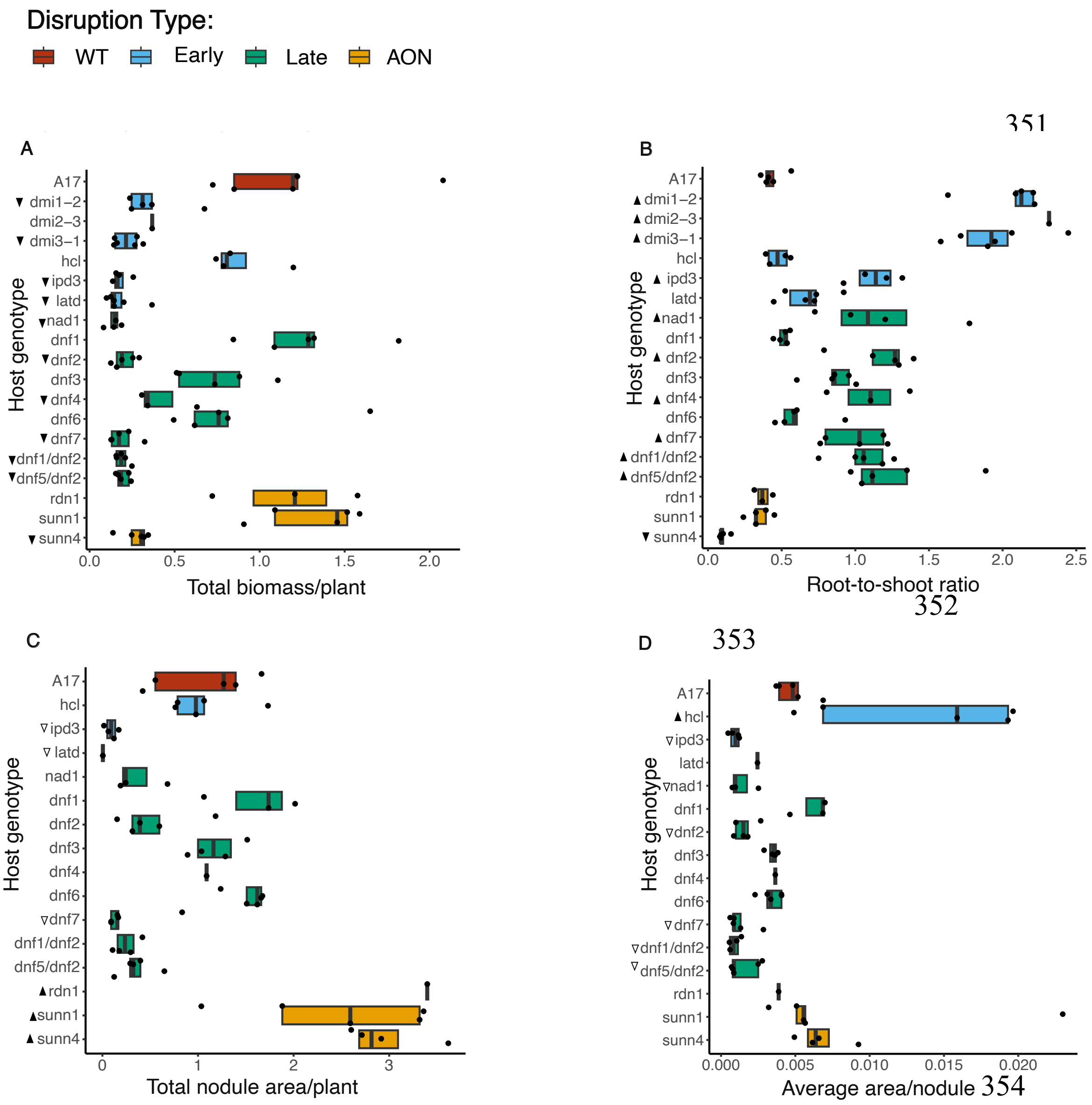

**Fig S8:** (A) Total biomass per plant and (B) root-to-shoot ratio were significantly altered in most host mutants compared to wild-type A17, with all but *sunn4* showing reduced biomass and increased root-to-shoot ratio. (C) Total nodule area per plant and (D) Average area per nodule represent host investment traits and were differentially affected across mutants. Triangles indicate significant differences relative to A17: upward arrow indicates increase; downward arrow indicates decrease. Boxplots are color-coded by the symbiotic stage disrupted in each host mutant. Closed triangles denote statistically significant differences relative to A17 (based on Tukey-adjusted *p* values). In contrast, open triangles denote statistically significant differences (based on Tukey-adjusted *P* values) relative to A17, as determined by $\log_{10}$-transformed data, which accounts for heteroscedasticity in trait variance.
